# CardioVinci: building blocks for virtual cardiac cells using deep learning

**DOI:** 10.1101/2021.08.22.457257

**Authors:** Afshin Khadangi, Thomas Boudier, Vijay Rajagopal

## Abstract

Recent advances in high-throughput microscopy imaging have made it easier to acquire large volumes of cell images. Thanks to electron microscopy (EM) imaging, they provide a high-resolution and sufficient field of view that suits imaging large cell types, including cardiomyocytes. A significant bottleneck with these large datasets is the time taken to collect, extract and statistically analyse 3D changes in cardiac ultrastructures. We address this bottleneck with CardioVinci.

---

Recent deep learning (DL) findings have paved the way for much accessible data quantification and analysis tools. These methods have shown unprecedented performance in a wide range of data analysis tasks, including image analysis. In biology, DL has improved the accuracy of segmenting EM data and the efficiency by minimising the time required for quantifying such datasets. Despite their successful applications in EM image analysis, these methods still represent some limitations, as reported in our previous works [1, 2]. Finding the exemplary DL architecture, optimising, and fine-tuning to segment various ultrastructures sometimes need sophisticated techniques, including ensemble learning. This makes it challenging to collect a large sample of cardiomyocytes, obtain optimal images from the samples, develop and fine-tune deep neural networks for segmenting organelles, and finally report statistical measures.

Generative adversarial networks (GANs) have been proposed to learn complex latent features from a dataset in an unsupervised manner. Simplistically speaking, GANs enable us to learn the original distributions of the data. For example, the EM images collected from dissected cardiac tissues only represent a sampled version from a complex data distribution representing the cardiomyocytes’ organisation within the tissue population. Many previous studies have utilised generative modelling to extract unsupervised feature representations across different cell types [3–8]. These studies are mainly carried out in 2D or are limited in resolution as one of the barriers to applications of GANs in biology used to be a low and poor resolution of projections of latent space.

Moreover, training high-resolution GANs in 3D are computationally exhaustive. However, the correlation among consecutive image slices in 3D microscopy image data can be utilised for passing 3D information during training 2D GANs. StyleGAN [9] addresses these challenges by adopting style transfer and stochastic variation in the generated images and demonstrably high-resolution, realistic images from latent space projections.

In this study, we propose CardioVinci, a designated workflow to statistically quantify 3D structures of mitochondria, myofibrils, and Z-disks without the sophisticated need for collecting large image samples of cardiomyocytes. Figure 1 represents the workflow of the proposed framework. CardioVinci is scalable and can be applied across other tissue types and image modalities. The proposed method enables the community to statistically quantify the properties of key organelles in cardiomyocytes, including spatial distributions, shape, and geometrical statistics. Moreover, we have proposed a novel training workflow that enables the users to generate realistic 3D reconstructions of cardiomyocyte architecture from 2D GANs. To the best of our knowledge, this is the first study that addresses 3D cell organisation using only 2D serial sections from focused ion-beam scanning electron microscopy (FIB-SEM) and using 2D segmentation and GAN setting. Moreover, this is the first study that reports high-resolution 3D virtual cardiac cell structure ranging from 10μm×10μm×2.5μm to 10μm×10μm×10μm.

**Figure 1.**
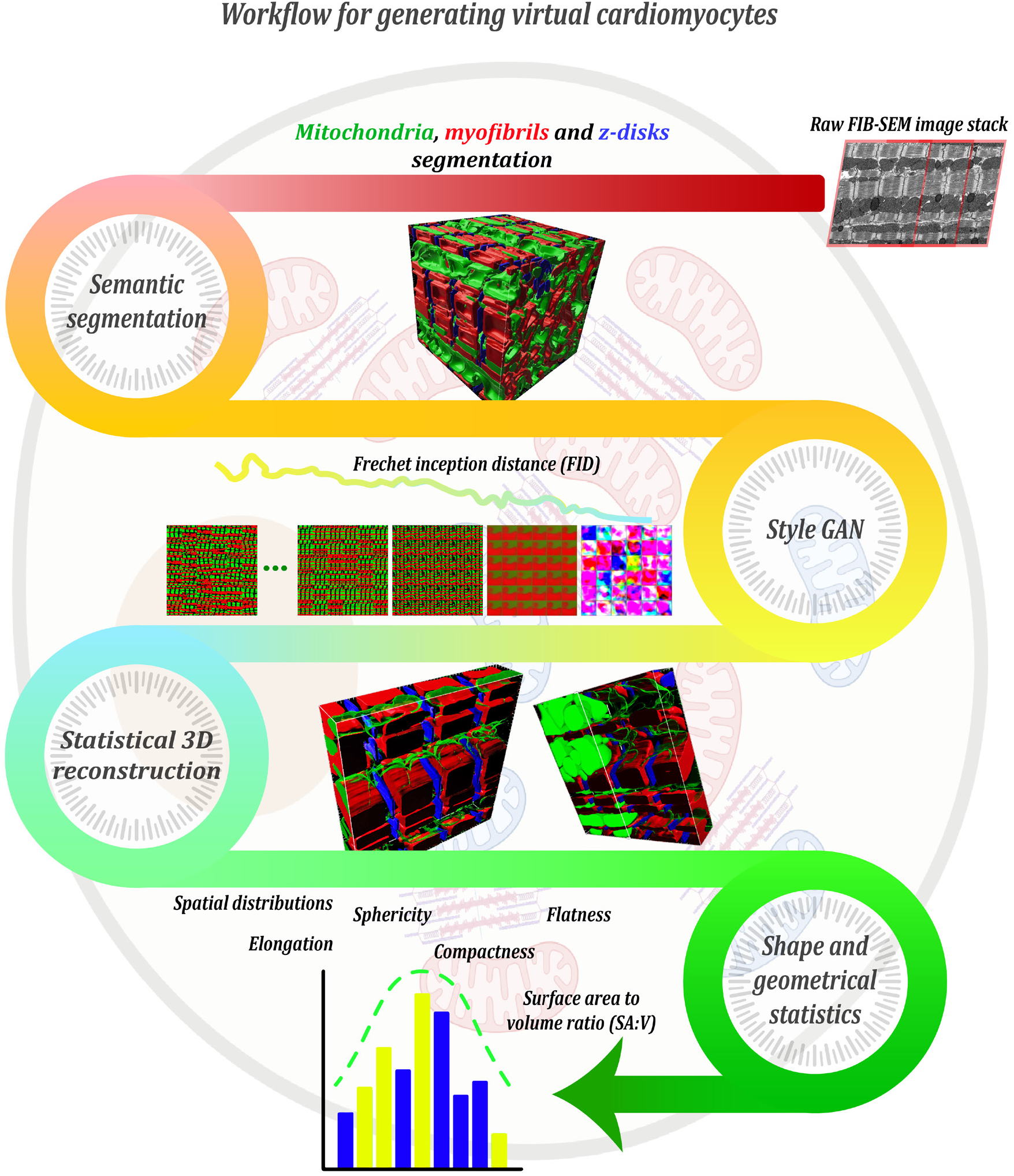
proposed pipeline for CardioVinci. First, semantic segmentation is used to segment target ultrastructures. Then StyleGAN is utilised to optimise the error between generator and discriminator. After the images are generated using the trained StyleGAN, 3D statistical representation of cardiomyocytes are reconstructed. And finally, 3D shape and geometric statistics are retrieved using the generated volume.

We used Style-GAN [9] in this study to extract key statistics of cardiomyocytes using generative modelling. Style-GAN is a 2D generative adversarial network that provides demonstrably distribution quality metrics. Despite its 2D architecture, we could translate the correlation across image stack slices into latent space by slightly adapting the training workflow. After we generated the random images using the optimised generator, we ordered the generated images using the Jaccard similarity score. Finally, we reconstructed the resulting image slices in 3D to visualise the statistical representation of cardiomyocytes using generative modelling. The reconstructed volumes are limited to 10μm×10μm×10μm and 10μm×10μm×2.5μm representing four sarcomeres as shown in Figure 2.

**Figure 2.**
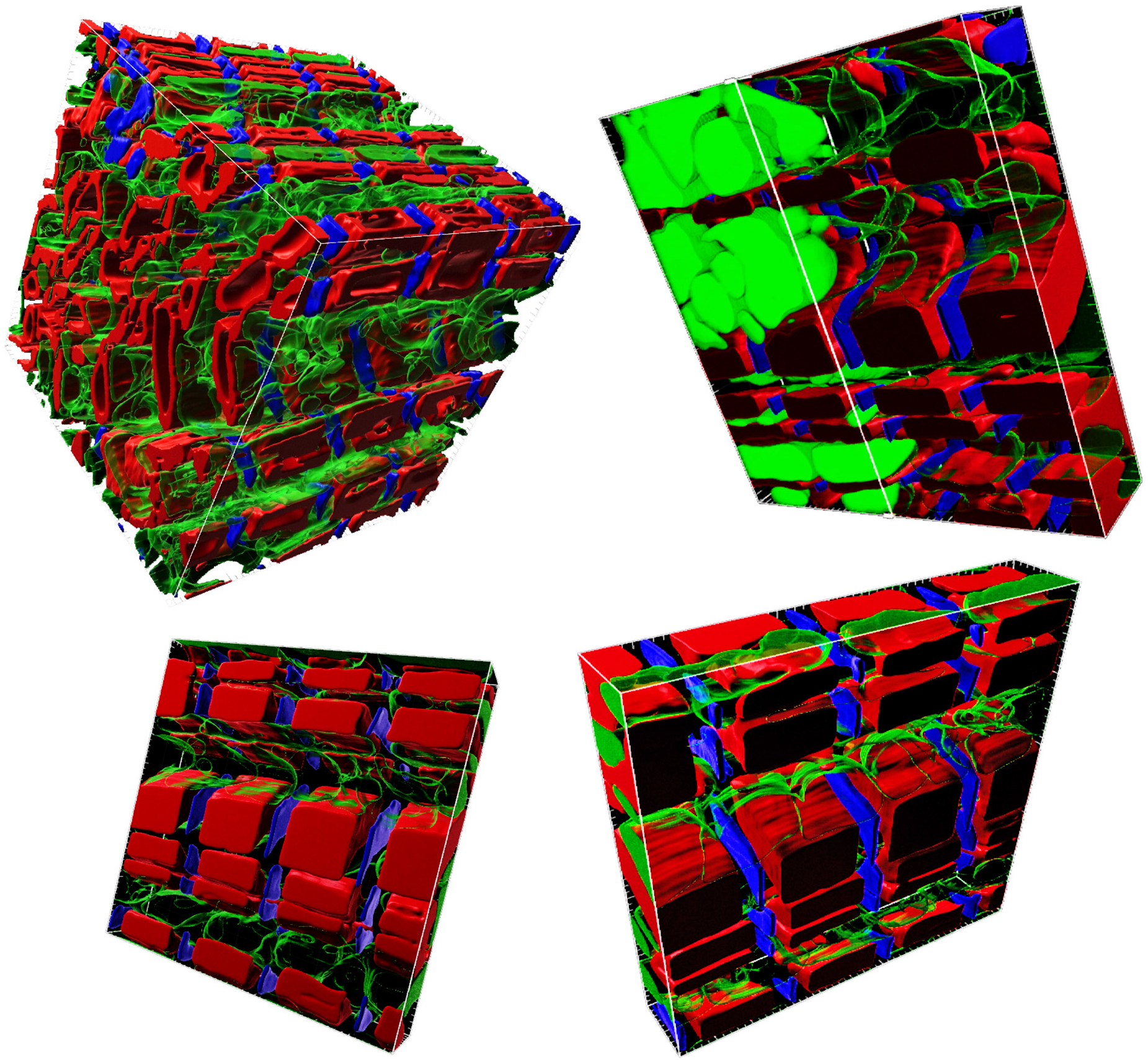
Results of the volume reconstructions from the GAN outputs. Red, blue and green represent the myofibrils, z-disks and mitochondria, respectively. All the volumes represent approximately four sarcomeres of the cardiomyocytes that have been generated using our trained GAN. The reconstructed volumes are limited to 10μm×10μm×10μm (top left) and 10μm×10μm×2.5μm.

The overarching goal of systems biology is to decode the complex ultrastructural dynamics within the cell to enable spatial simulation of such functional subcellular networks [10]. Generative models allow the creation of a spatial organisation of the cells directly from the microscopy images. These models encode the statistical variation across the cell architecture and capture the shape, size and spatial distribution of subcellular ultrastructures. CardioVinci captures the spatial organisation of the cardiomyocytes directly from the FIB-SEM and SBF-SEM images. The outputs of the CardioVinci can be used for developing biochemical models of the cardiomyocytes as it provides statistically accurate spatial organisation. In this study, we extracted relative spatial distributions for mitochondria, myofibrils, and Z-disks using 3D point statistics based on the results of GAN (supplementary).

In addition to the above, we extracted surface area to volume ratio (SA:V) along with other 3D shape and geometrical statistics, including elongation, compactness, flatness, spareness and sphericity. These statistics are drawn based on the results of GAN and segmented images from the original dataset. The results suggest that GAN and segmented images share similar mean and median across these statistics; however, we observe relatively higher variance across extracted distributions using GAN. Our analysis suggests that Z-disks represent higher mean, median, and variance for SA:V, whereas mitochondria demonstrate a lower mean and variance range for SA:V. The latter applies to elongation, too, as Z-disks show a higher mean, median, and variance range for elongation, whereas mitochondria demonstrate lower values. Thus, compactness is minimal for Z-disks and maximal for mitochondria. Moreover, mitochondria and Z-disks show relatively the same statistics for flatness, whereas myofibrils show a lower mean, median, and variance range for flatness. In addition to the above, mitochondria and myofibrils represent relatively similar statistics for spareness and sphericity. However, Z-disks possess minimal sphericity properties.

We used CardioVinci to analyse mitochondria and myofibrils shape and geometric statistics in type 1 diabetic cardiomyocytes. In both experiments, we have used generative modelling to obtain the statistics as highlighted above. Our results show that SA:V has been reduced in the diabetic sample compared to the control sample (mitochondrial SA:V has been altered drastically in the diabetic sample). In addition, mitochondria and myofibrils show increased compactness and sphericity in the diabetic condition compared to the control. These findings align with the previous results on mitochondrial properties on diabetic induced cardiomyocytes using 2D analysis on 2D data [11]. We found a considerable difference between control and diabetic samples shape and geometric statistics, especially in mitochondria.

CardioVinci is a framework to reconstruct the statistical representation of cardiomyocytes in 3D. Although many studies have addressed biological questions using generative adversarial networks [3–5, 7, 12–16], most of these studies report their findings in 2D or are limited in resolution due to the image modalities. Moreover, 3D generative modelling has not been conducted for quantifying scanning electron microscopy data to date. In this paper, we have proposed to segment crucial ultrastructures in cardiomyocytes semantically. Then the segmentation results are used to train the Style-GAN to learn the corresponding statistical distributions in a generative manner. We have also extended our study to analyse the changes in the mitochondrial and myofibrillar statistics in the presence of diseases such as diabetes. We have modified the Style-GAN in a way that is able to learn the data correlations between consecutive image slices. Hence, this will be the first study that reports 3D cardiomyocyte geometric statistics using a 2D GAN setting.

To summarise, CardioVinci is a scalable framework that can generate the statistical representations of electron microscopy data. CardioVinci exhibits phenomenal performance in the presence of limited data due to its scalability and would represent optimal performance when a large dataset is available. Our future aim is to utilise the style mixing capability of the StyleGAN to generate synthesized variations across different tissue blocks or even tissue types. We hope CardioVinci be utilised across other tissue types, and of course, other image modalities. We suggest the readers refer to the supplementary text and figures for detailed methods and analysis.

## Supplementary text

### Data

We used one publicly available FIB-SEM cardiac dataset in this study, as highlighted in [1]. The second dataset includes left ventricular myocyte serial block-face scanning electron microscopy (SBF-SEM) image datasets collected from type 1 diabetic induced mice, as described previously [11]. We extracted 30 random patches from each dataset, each having 512 × 512 pixels for training, testing and validation. After we manually annotated mitochondria, myofibrils and Z-disks on the first sample and mitochondria, myofibrils on the second sample. We split data randomly into training, validation and testing by 20/30, 5/30, and 5/30, respectively. All the random data splits were performed using K-fold cross-validation, and the inference performance is reported based on the best fold model.

### Training and testing

All the experiments in this study were implemented using Deep Learning AMI (Amazon Linux 2) Version 46.0 using Amazon Web Services (AWS). These experiments were performed on GPU instances. We used “g4dn.12xlarge” (4 GPUs of a total of 64 GiB memory) and “g4dn.metal” (8 GPUs of a total of 128 GiB memory) for segmentation and training the GAN, respectively.

### Semantic segmentation improves feasibility and accuracy

Machine learning (ML) and DL methods offer a fast and accurate segmentation of various organelles. However, segmenting some of the ultrastructures such as Z-disks has its own challenges using ML or DL methods as shown in [17, 18]. This is mainly due to a relatively large label imbalance. For example, when the aim is to segment Z-disks only (binary segmentation), a relatively large label imbalance leads to poor optimisation performance. However, in this work, we show that segmenting Z-disks along with mitochondria and myofibrils not only boosts the feasibility of Z-disks segmentation using DL methods but also leads to higher test data performance across various metrics, including *V^Rand^*_(*thinned*)_ and *V^Info^*_(*thinned*)_. We modified U-net [19] architecture using a fully connected bottleneck and added a softmax layer as the output node for segmenting three ultrastructures of our interest. More details can be found in [2]. Figure 1 represents evaluation metrics for mitochondria, myofibrils, and Z-disks on the test sample.

### Spatial distributions reveal the dominance of clustered patterns in cardiomyocytes

We used GAN results to extract relative spatial distributions for mitochondria, myofibrils, and Z-disks using 3D point statistic. Our results show that mitochondria are distributed in a clustered pattern relative to myofibrils up to between 60 nm to 70 nm. As we move further away, the mitochondria distributions become uniform. The results also show that mitochondria are distributed in a clustered pattern relative to Z-disks up to 70 nm. In addition to the above, the results also show that myofibrils are distributed in a clustered pattern relative to mitochondria and Z-disks up to 70 nm; however, they represent a relatively uniform pattern beyond 70 nm. And finally, our results show that Z-disks demonstrate dominantly clustered patterns up to 80 nm relative to mitochondria and myofibrils, whereas they show uniform patterns beyond that point. Figure 2 shows the spatial distributions.

### StyleGAN can be used to learn correlations across 3D microscopy image slices

Thanks to the TFRecord formatting, we can store a set of features along with the image itself in the TFRecord data. We used one hot-encoded formatting to assign a label to two consecutive image slices as part of building our TFRecord data for training the StyleGAN using segmented image slices. Then we added another input node to the StyleGAN as an embedding layer to embed the correlation across different slices. We used fully connected layers to capture the nonlinearity across image slices. We followed exactly the same strategy to optimise StyleGAN as highlighted in [9], except that we did employ shuffling during the training. After the network had been converged based on Fréchet inception distance (FID), we generated the images using the generator. We then ordered the resulted images according to the minimised Jaccard distance between the generated slices to obtain 3D virtual volume of cardiomyocytes.

### 3D statistics reveal increased SA:V for the cardiomyocytes in the presence of diabetes

In the previous study [11], the authors have suggested increased SA:V for mitochondrial clusters. The authors have utilised cluster perimeters to estimate SA:V in 2D for the mitochondrial clusters. However, based on 3D statistics, we found that cardiomyocytes exhibit decreased SA:V in the presence of diabetes. Our understanding is that mitochondrial fusion and fission represent sophisticated dynamics from the morphological perspective; hence, the 2D analysis should not be conclusive. We found that despite transversal mitochondrial fission in the diabetic cardiomyocytes, they represent longitudinal fusion with other mitochondria clusters, which leads to increased mitochondrial volume. Figure 4 represents the statistical measurements based on 3D shape and geometric statistics for control and diabetic cardiomyocytes.

## Supplementary figures and tables

**Figure 1.**
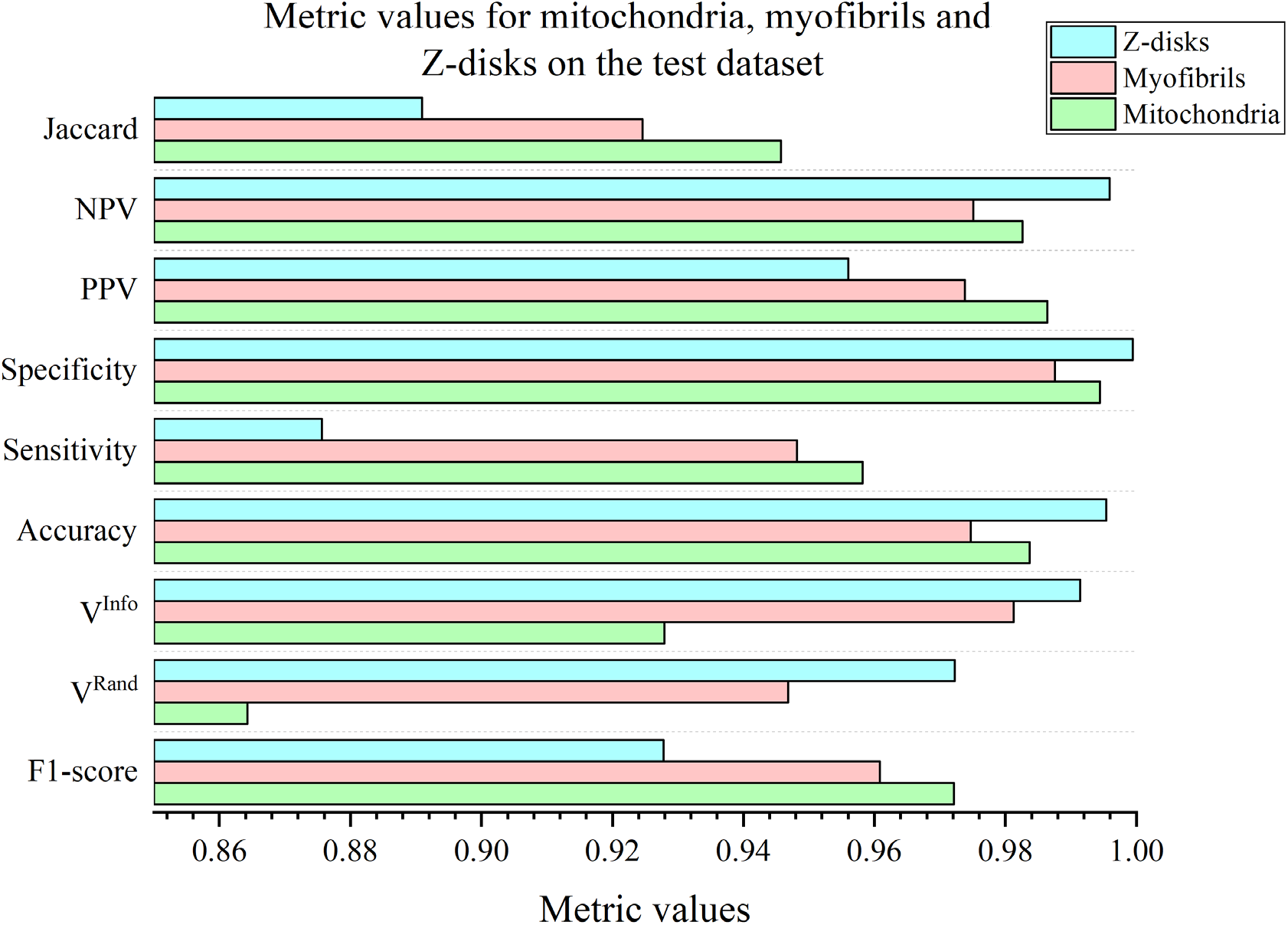
Evaluation metrics for the segmentation results based on test data. Z-disks have achieved the maximum *V^Rand^*_(*thinned*)_, *V^Info^*_(*thinned*)_, accuracy, specificity and NPV scores compared to mitochondria and myofibrils. PPV and NPV represent positive predicitve and negative predictive values, respectively.

**Figure 2.**
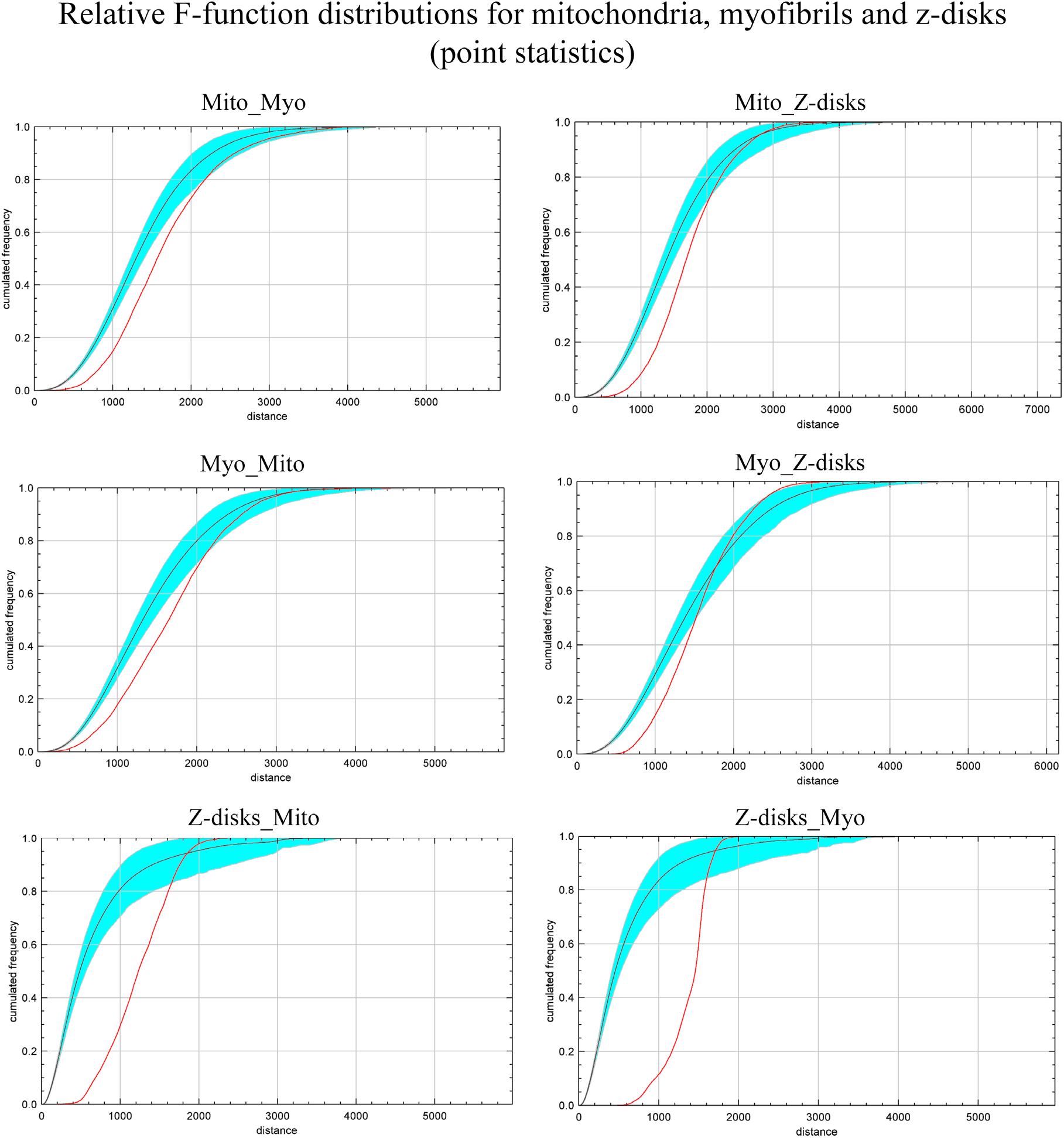
Cumulative F-function distributions for 3D point statistics. Mito and Myo represent mitochondria and myofibrils, respectively. Each of the individual subplots represent the F-function for the left organelle relative to the right one. For example, the top left subplot “Mito_Myo” represents the F-function for mitochondria relative to the myofibrils. Shaded area (cyan) represent the envelopes, black and red lines represent theoretical and observed values, respectively. Distance is in nm.

**Figure 3.**
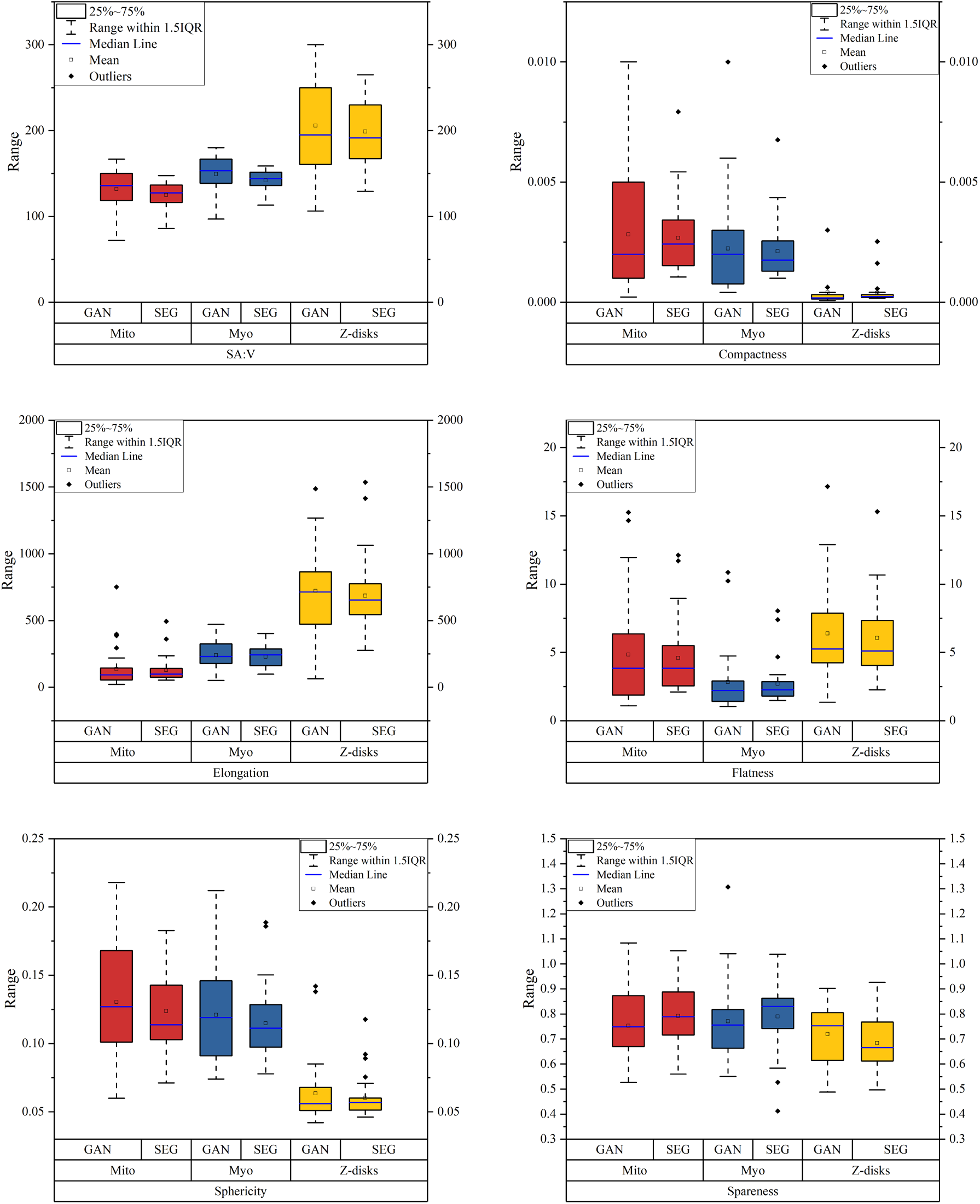
Surface area to volume ratio (SA:V) along with other 3D shape and geometrical statistics including elongation, compactness, flatness, spareness and sphericity. These statistics are drawn based on the GAN results and segmented original dataset (SEG).

**Figure 4.**
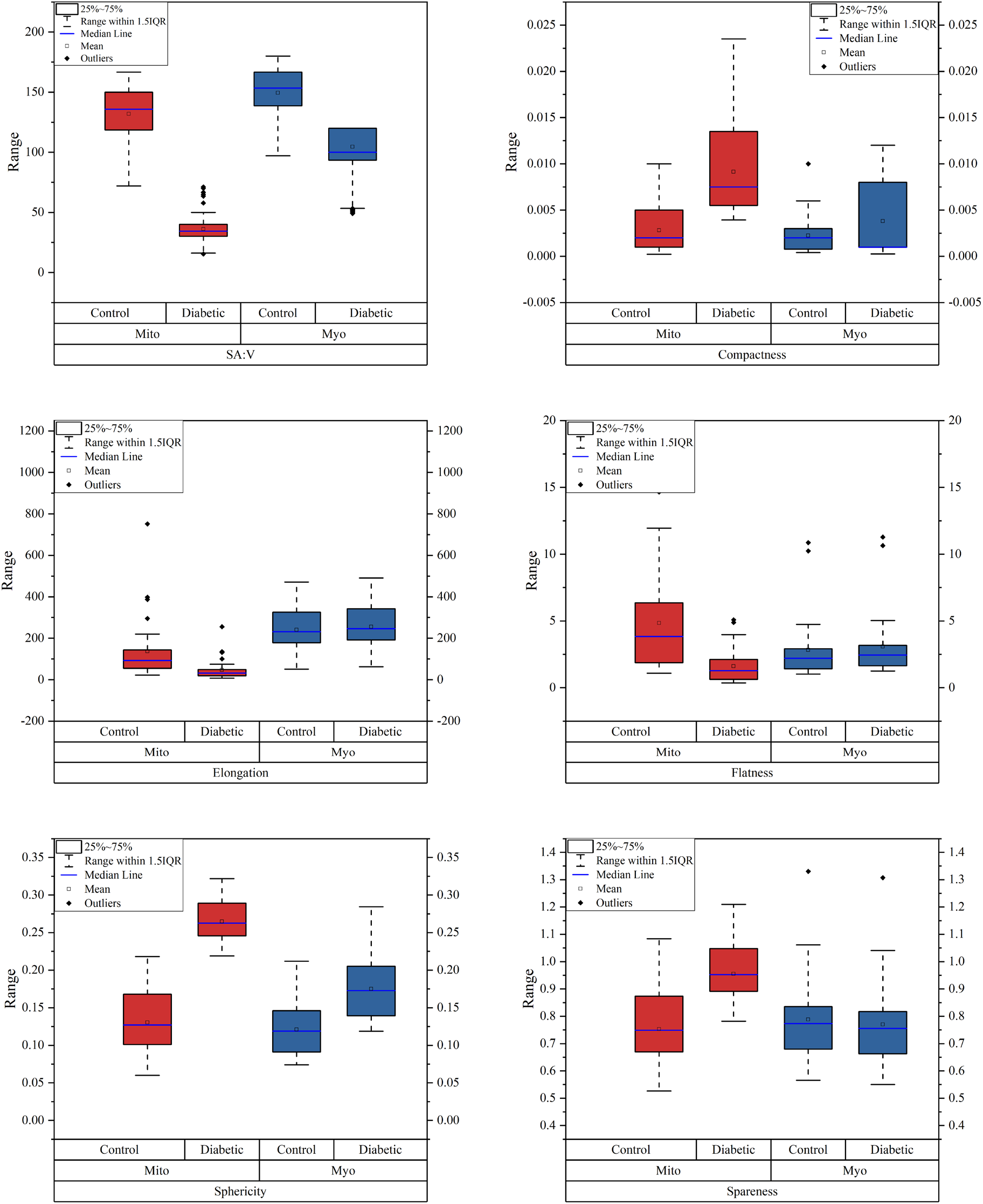
Surface area to volume ratio (SA:V) and other 3D shape and geometrical statistics, including elongation, compactness, flatness, spareness, and sphericity. These statistics are drawn based on the GAN results for both control and diabetic samples.

**Figure 5.**
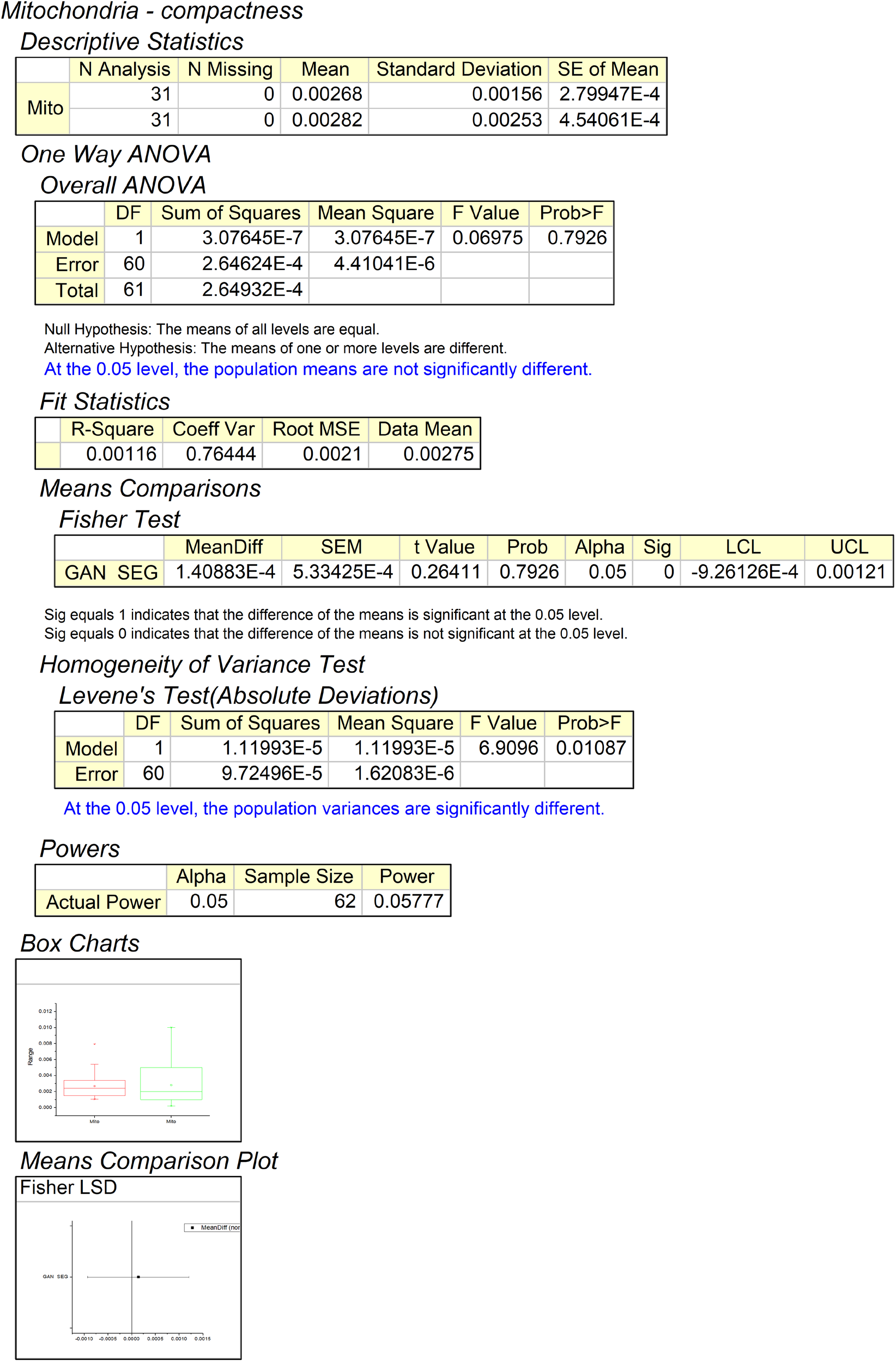
Hypothesis testing using one way ANOVA for mitochondria compactness significance between GAN outputs and segmentation results using control dataset.

**Figure 6.**
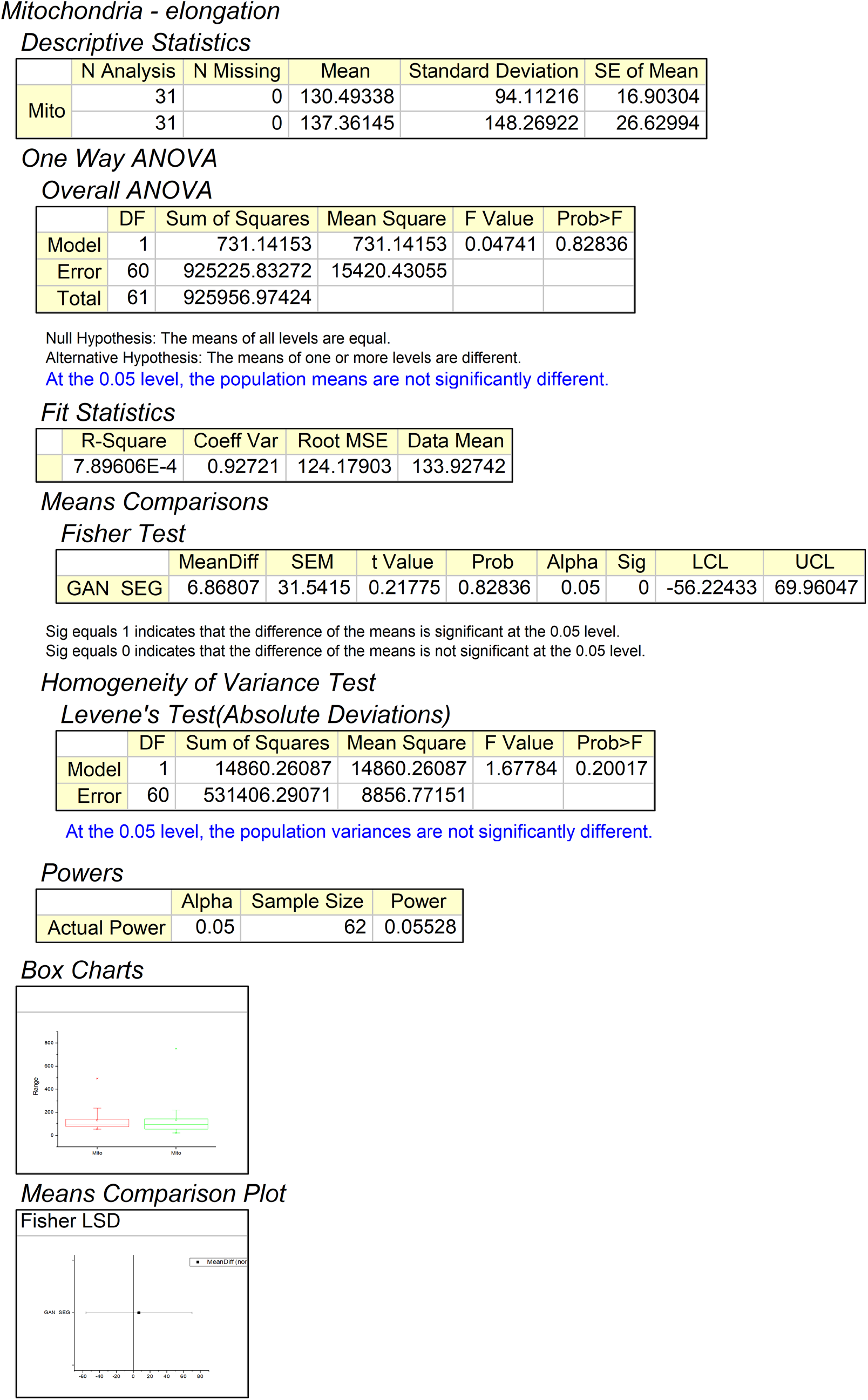
Hypothesis testing using one way ANOVA for mitochondria elongation significance between GAN outputs and segmentation results using control dataset.

**Figure 7.**
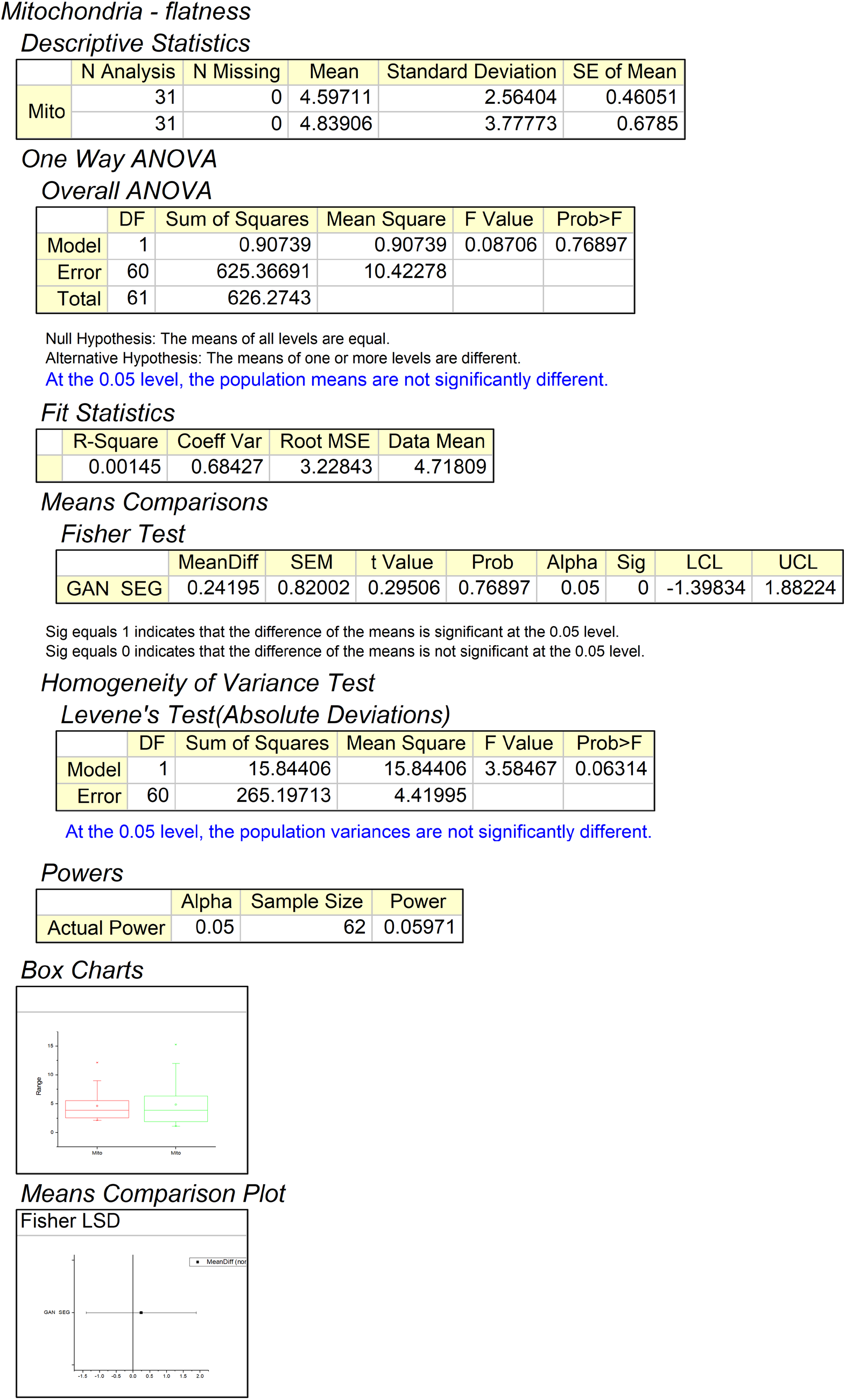
Hypothesis testing using one way ANOVA for mitochondria flatness significance between GAN outputs and segmentation results using control dataset.

**Figure 8.**
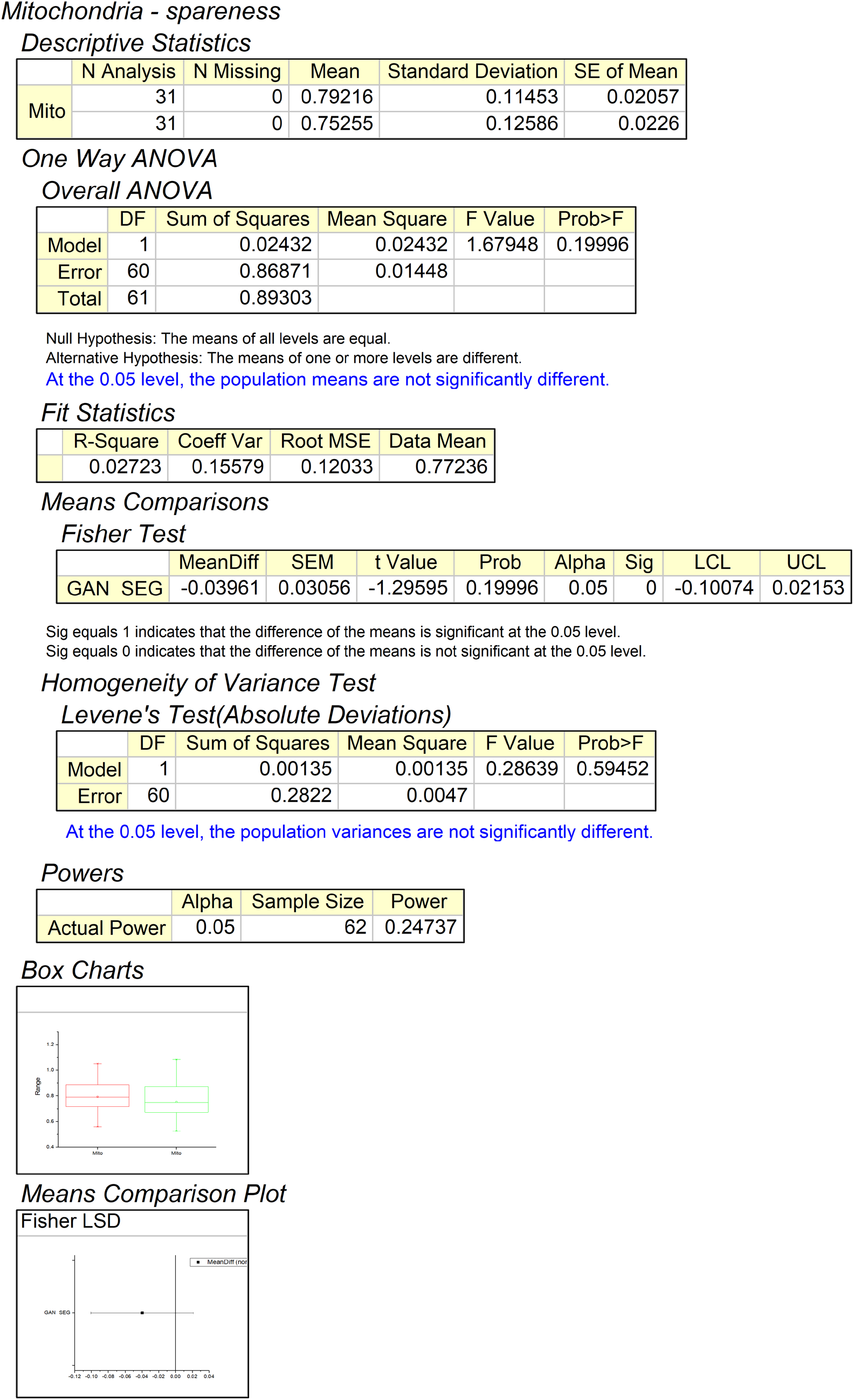
Hypothesis testing using one way ANOVA for mitochondria spareness significance between GAN outputs and segmentation results using control dataset.

**Figure 9.**
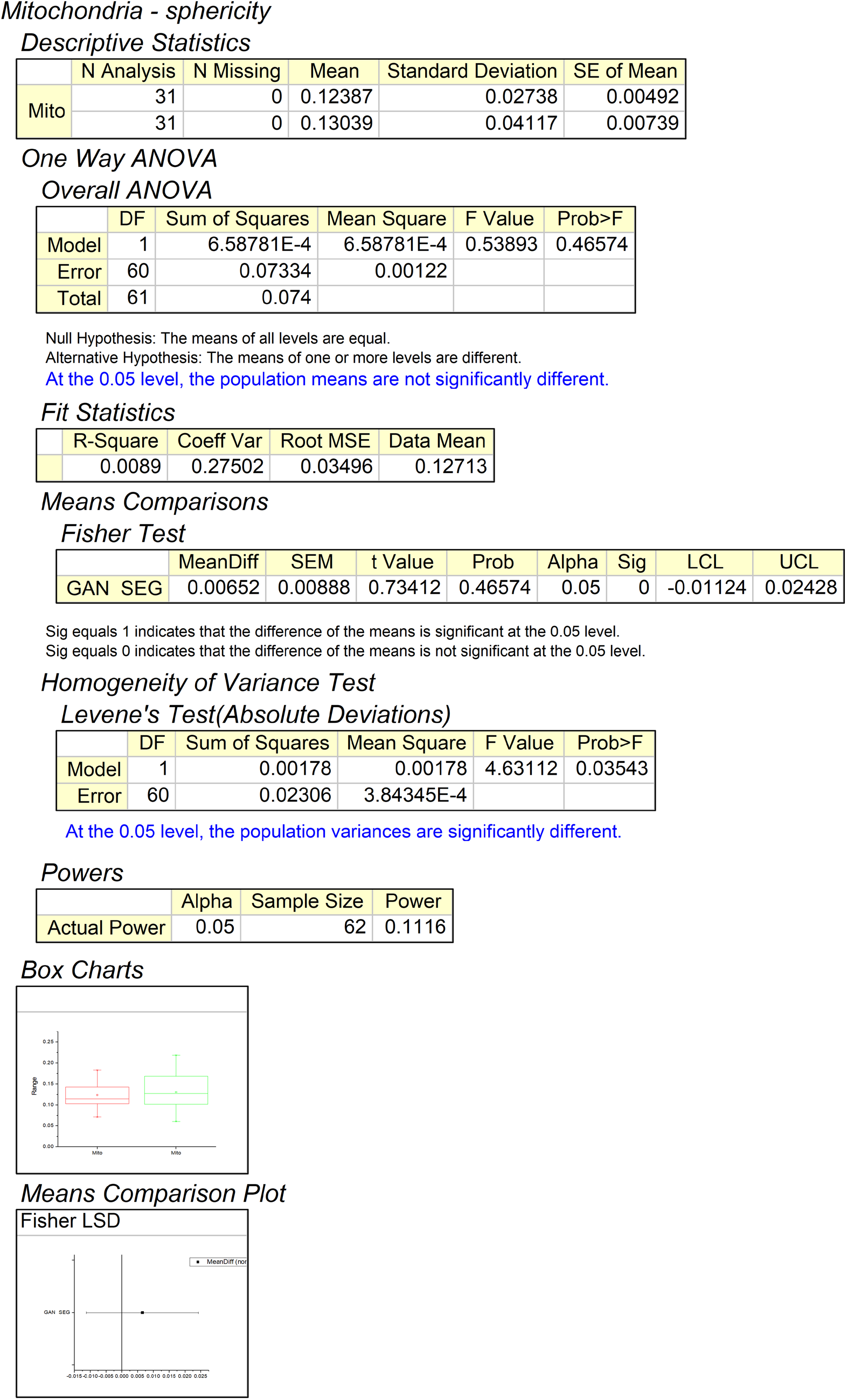
Hypothesis testing using one way ANOVA for mitochondria sphericity significance between GAN outputs and segmentation results using control dataset.

**Figure 10.**
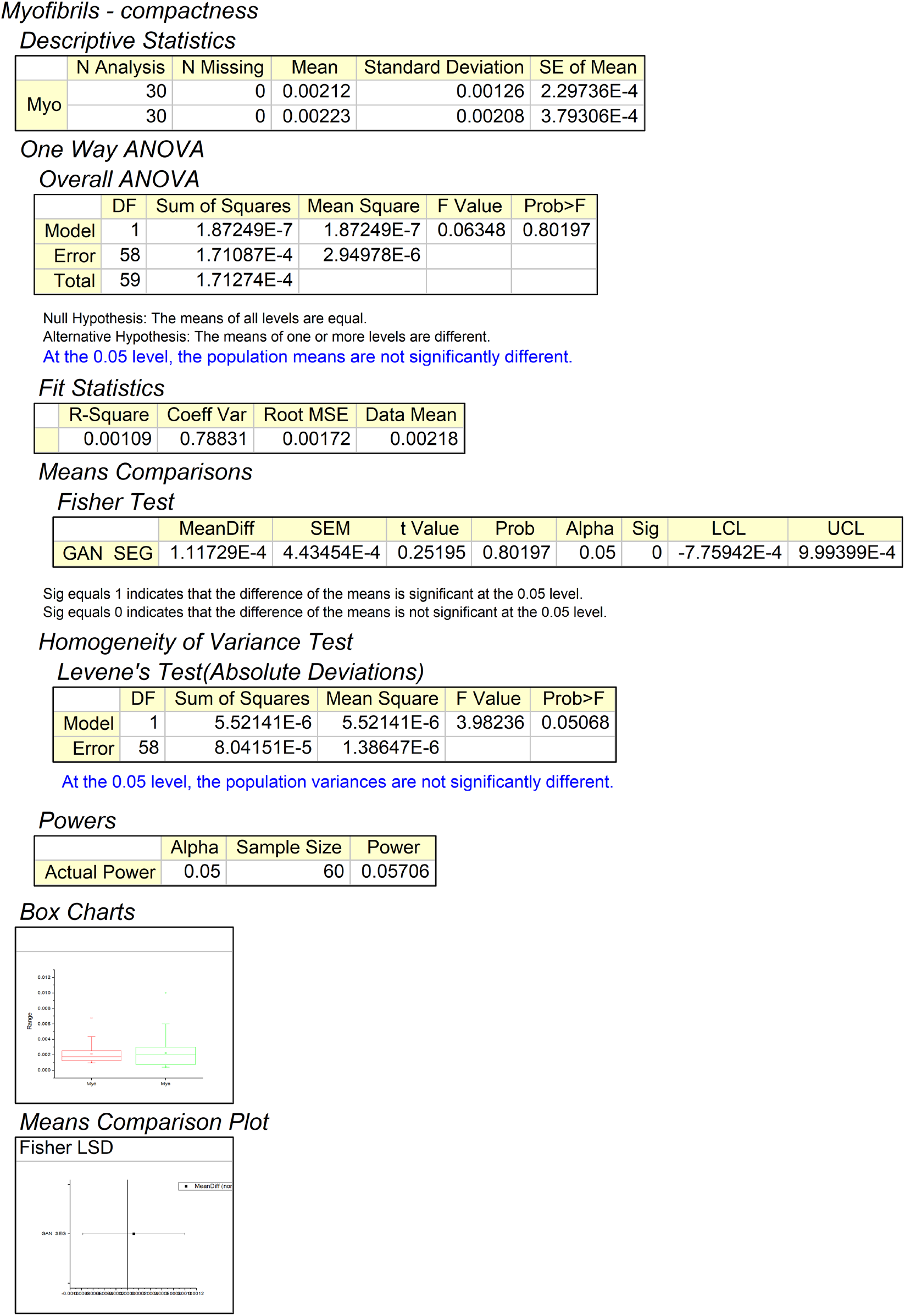
Hypothesis testing using one way ANOVA for myofibrils compactness significance between GAN outputs and segmentation results using control dataset.

**Figure 11.**
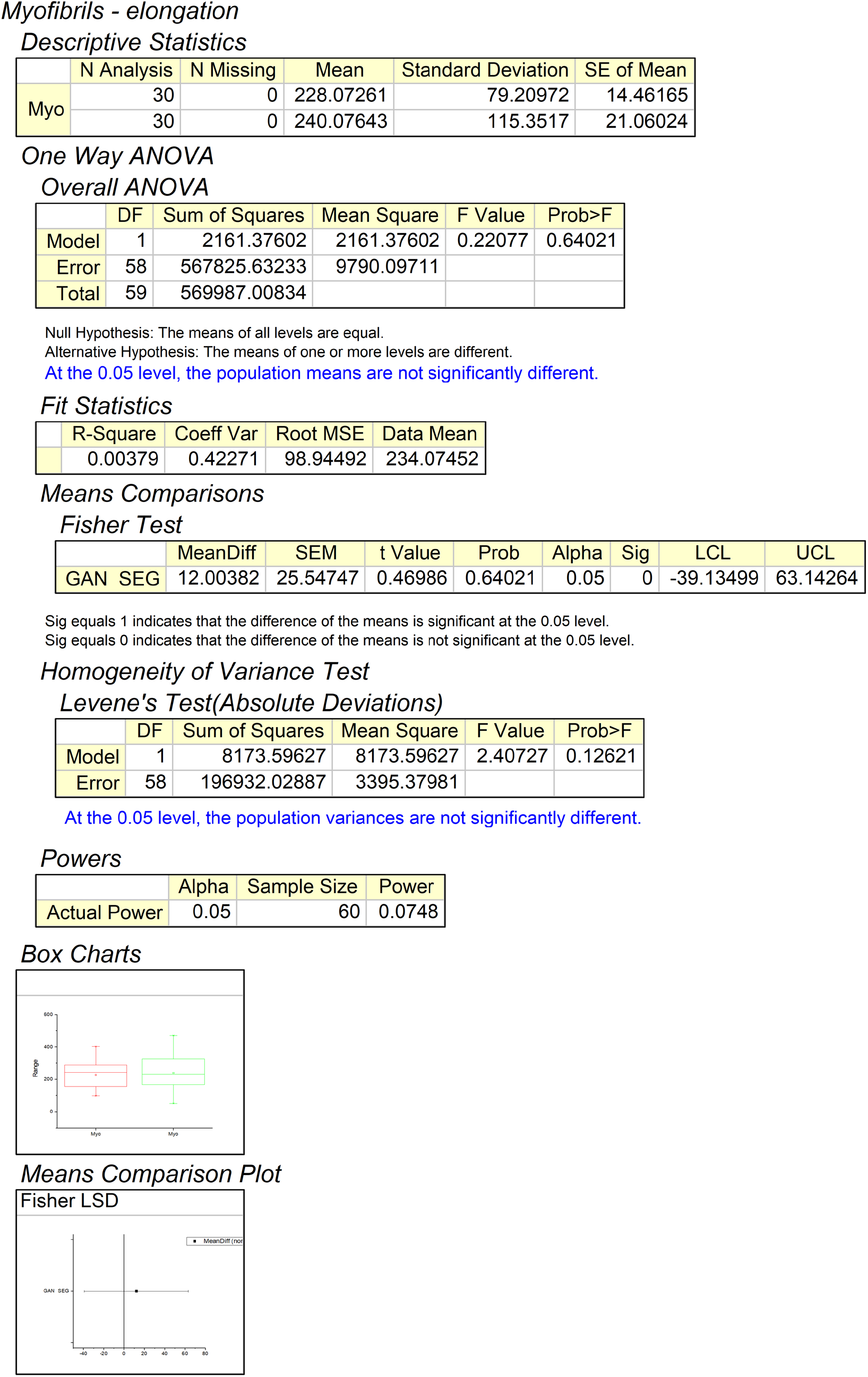
Hypothesis testing using one way ANOVA for myofibrils elongation significance between GAN outputs and segmentation results using control dataset.

**Figure 12.**
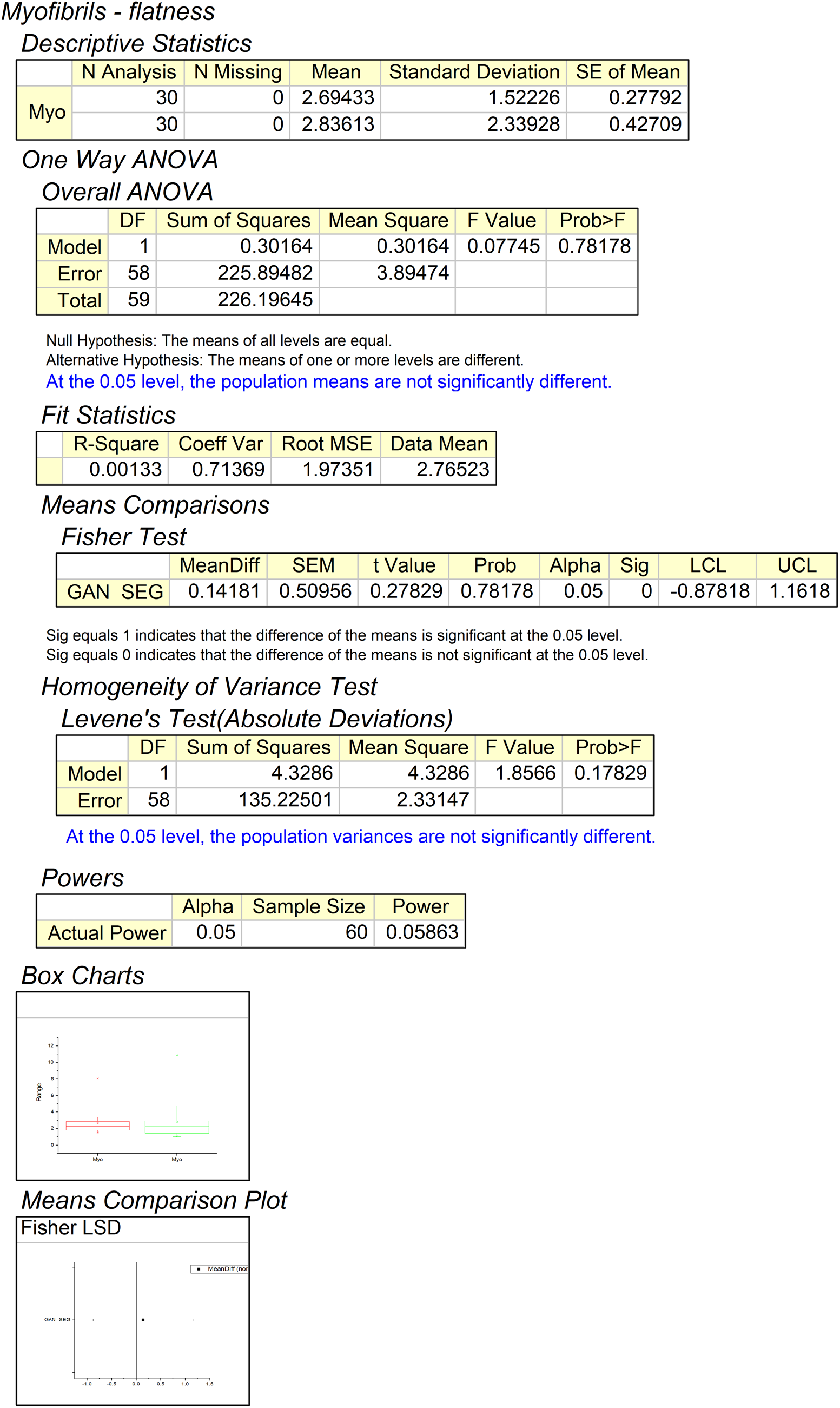
Hypothesis testing using one way ANOVA for myofibrils flatness significance between GAN outputs and segmentation results using control dataset.

**Figure 13.**
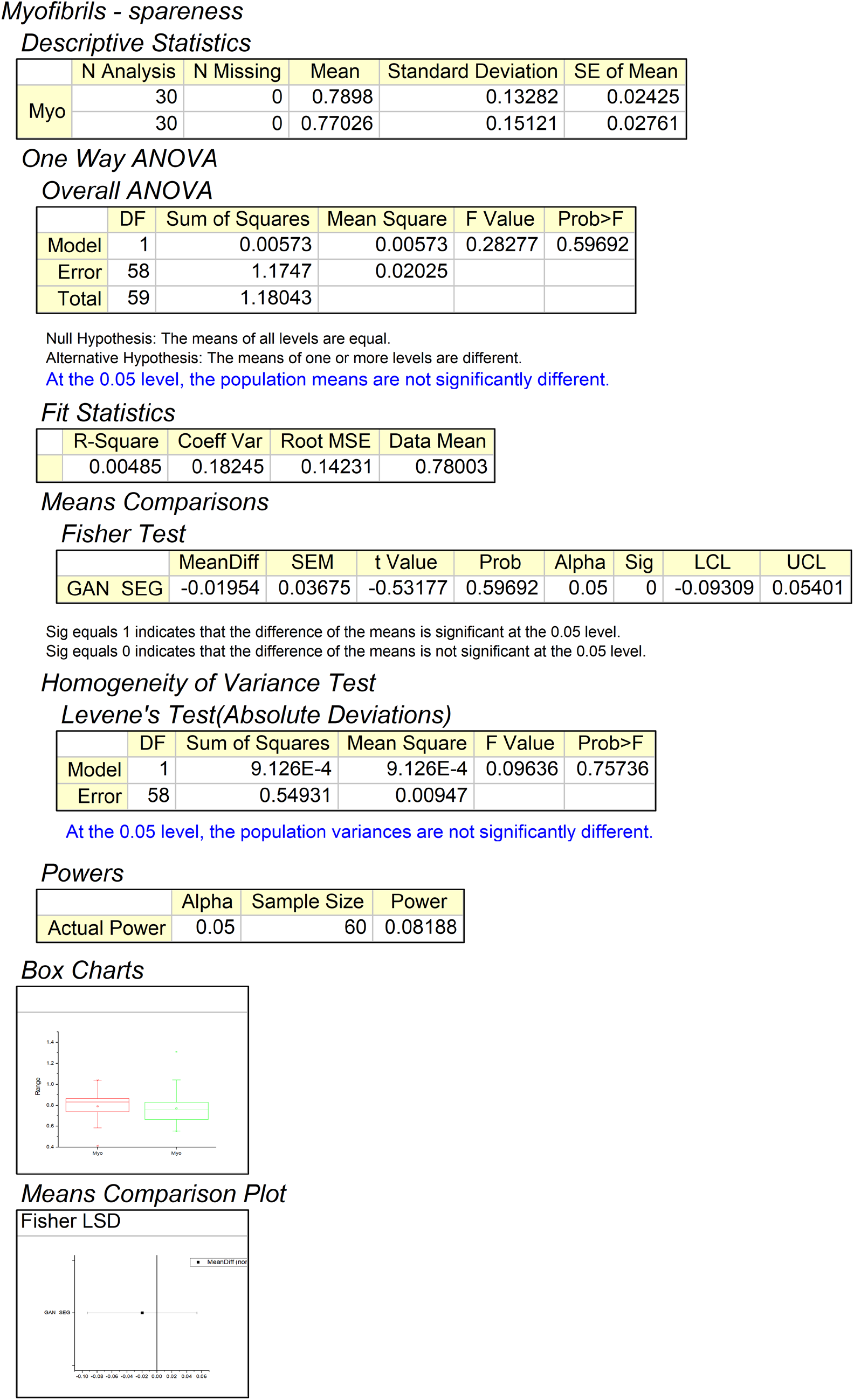
Hypothesis testing using one way ANOVA for myofibrils spareness significance between GAN outputs and segmentation results using control dataset.

**Figure 14.**
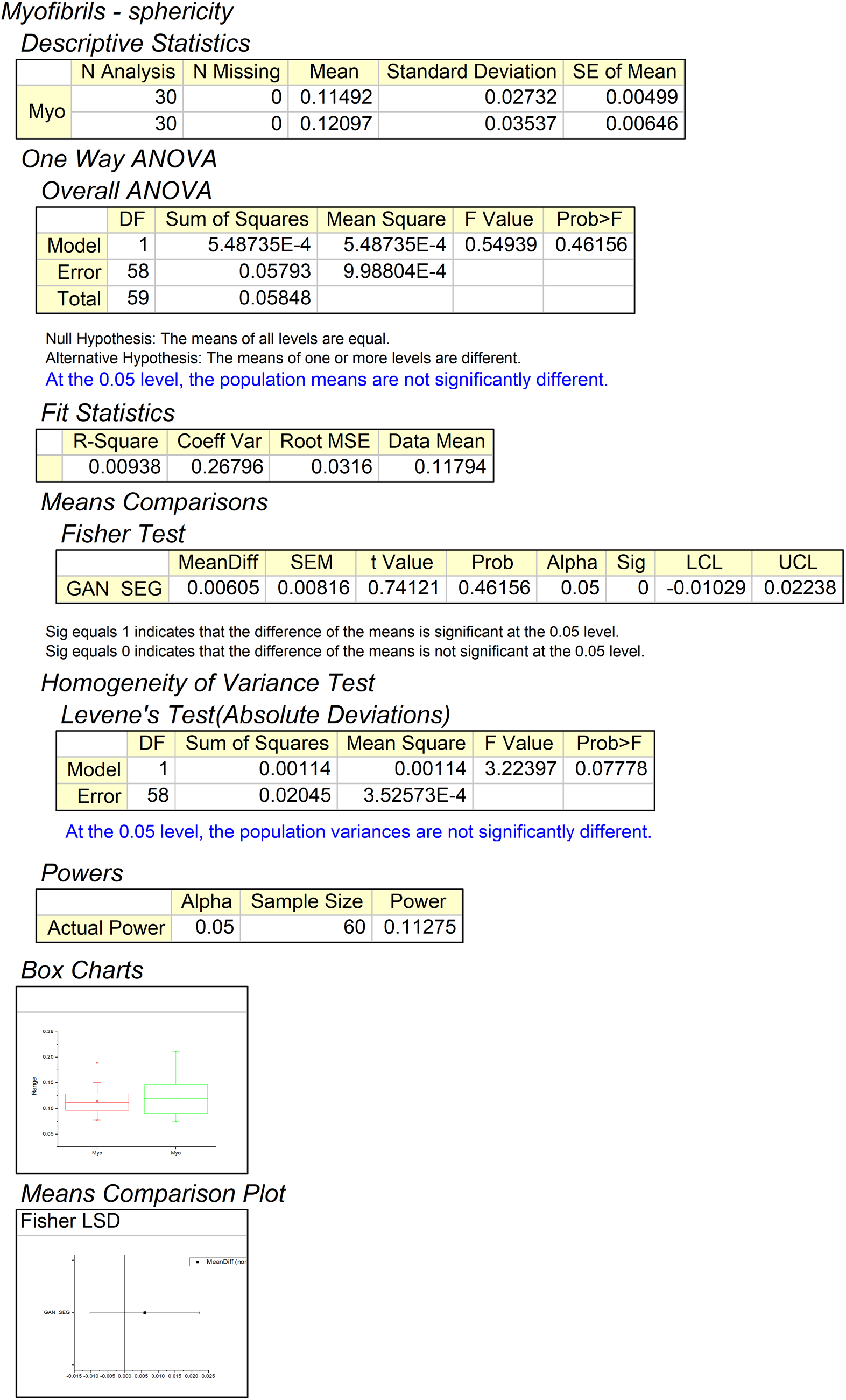
Hypothesis testing using one way ANOVA for myofibrils sphericity significance between GAN outputs and segmentation results using control dataset.

**Figure 15.**
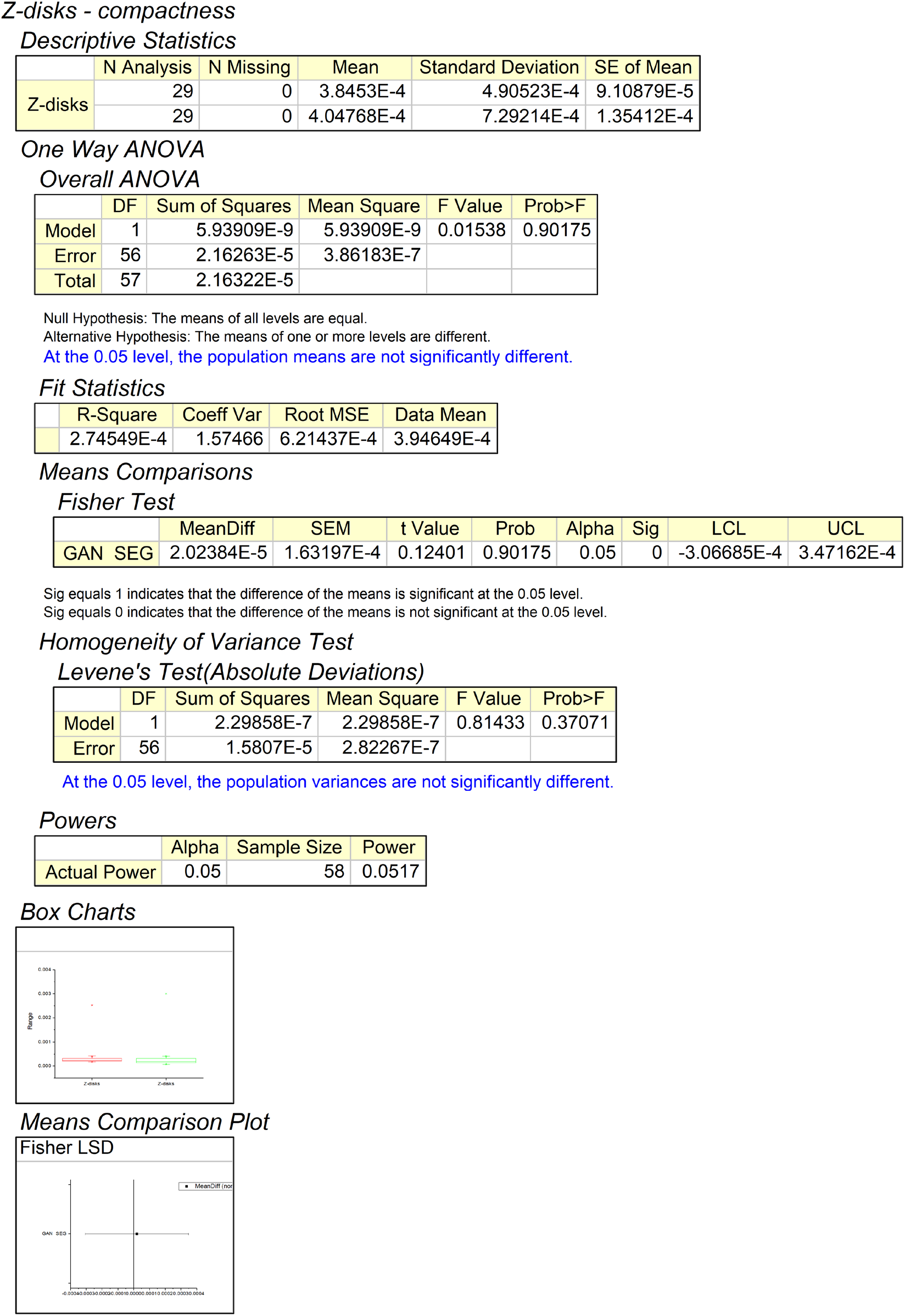
Hypothesis testing using one way ANOVA for z-disks compactness significance between GAN outputs and segmentation results using control dataset.

**Figure 16.**
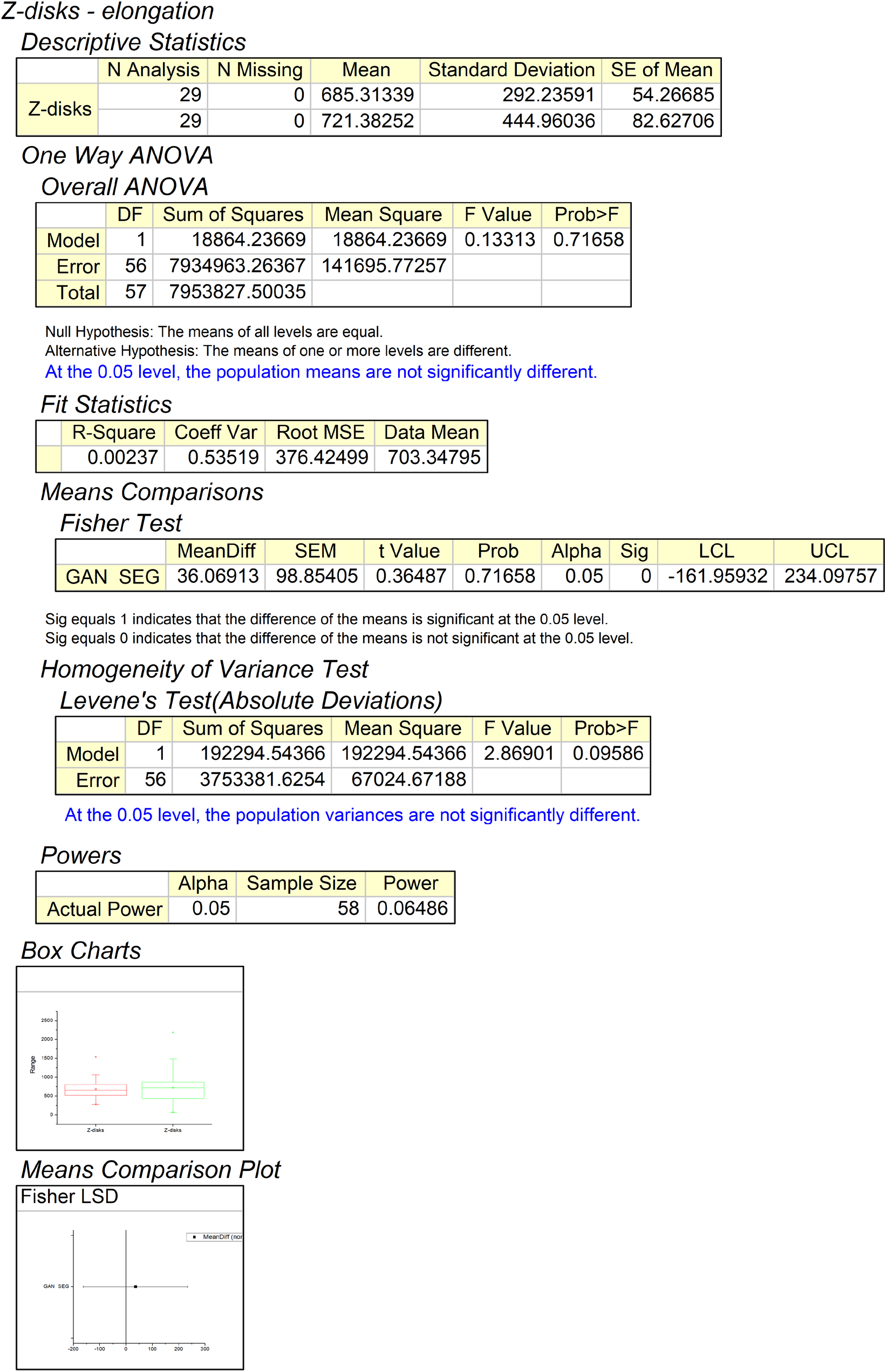
Hypothesis testing using one way ANOVA for z-disks elongation significance between GAN outputs and segmentation results using control dataset.

**Figure 17.**
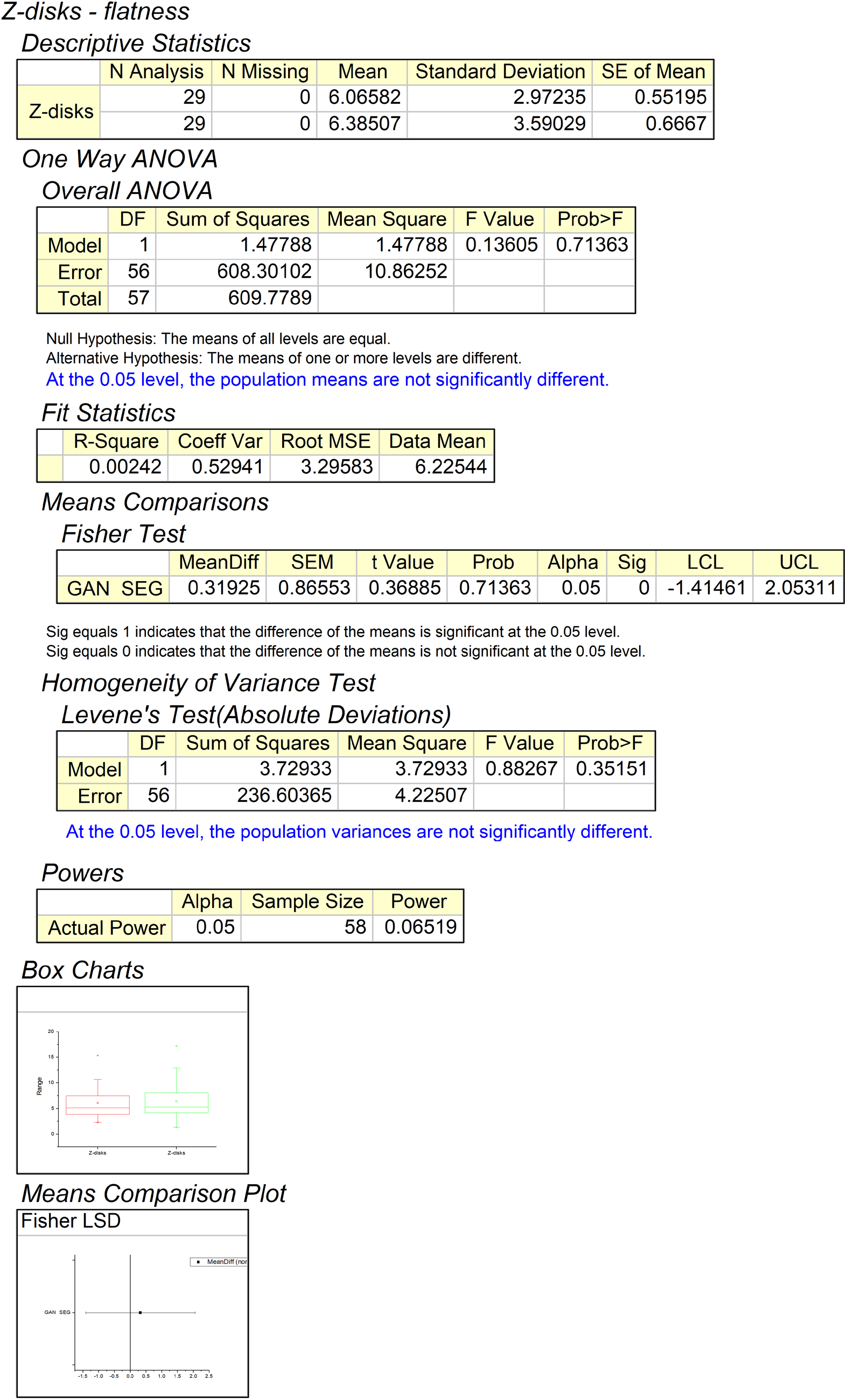
Hypothesis testing using one way ANOVA for z-disks flatness significance between GAN outputs and segmentation results using control dataset.

**Figure 18.**
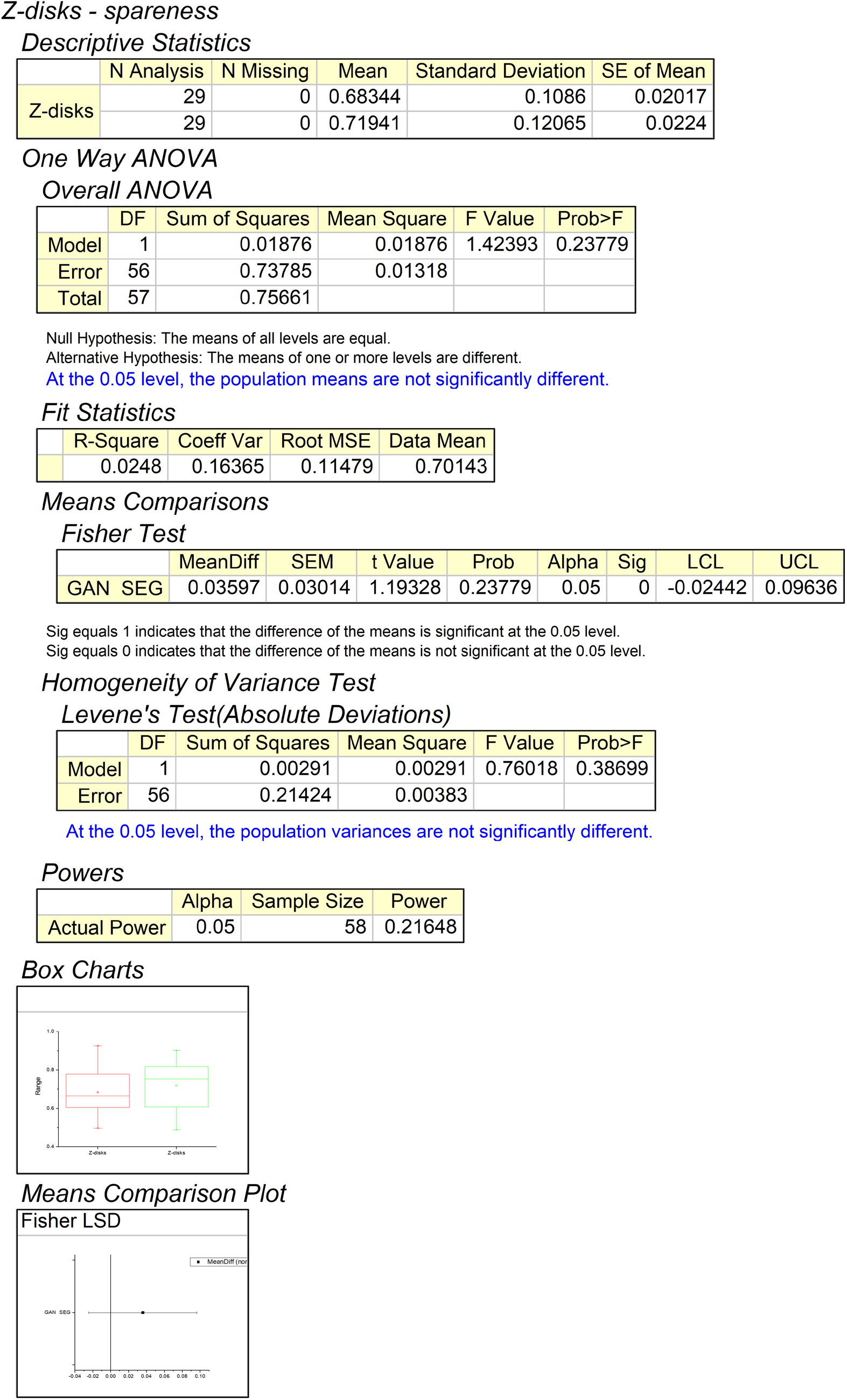
Hypothesis testing using one way ANOVA for z-disks spareness significance between GAN outputs and segmentation results using control dataset.

**Figure 19.**
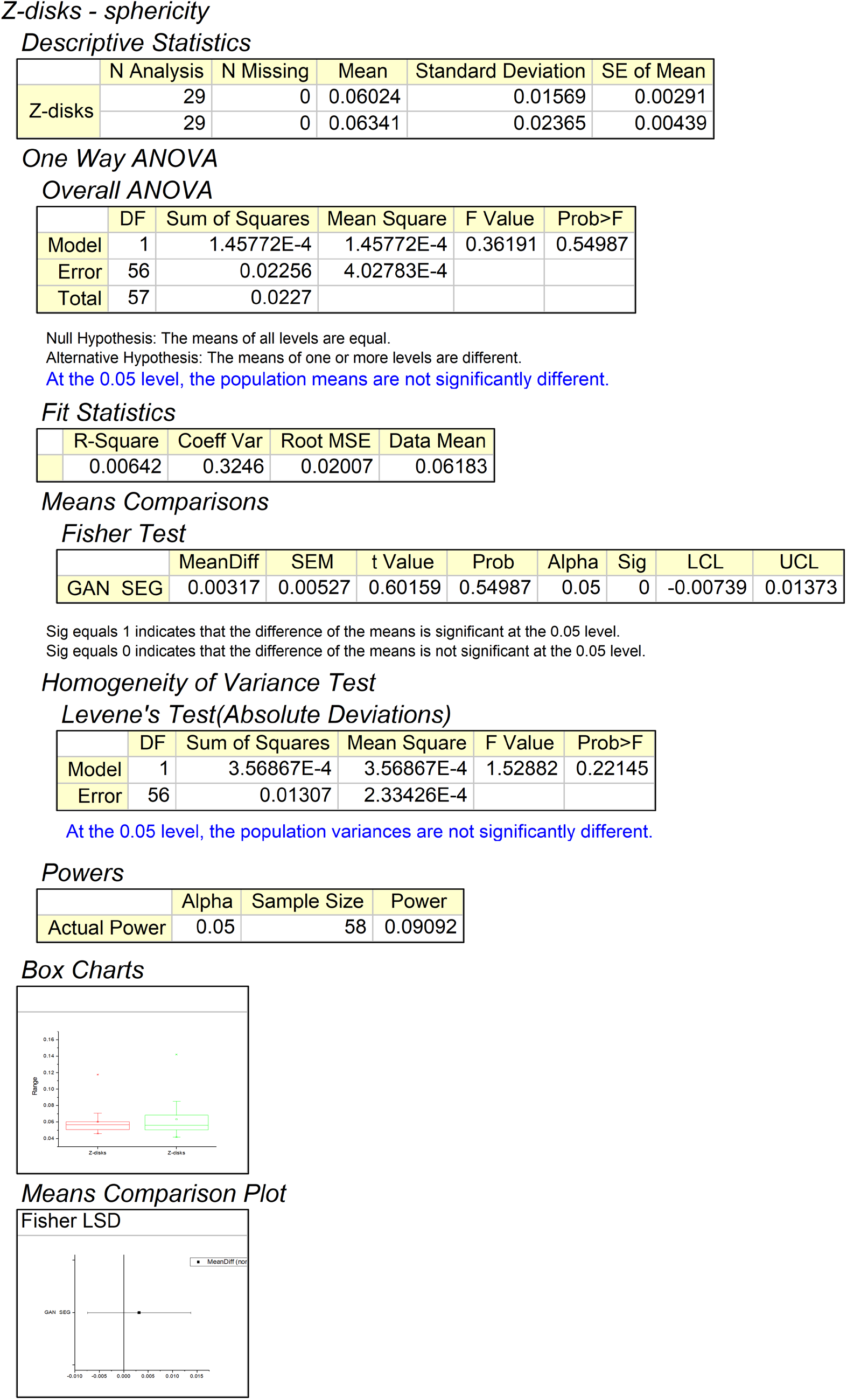
Hypothesis testing using one way ANOVA for z-disks sphericity significance between GAN outputs and segmentation results using control dataset.

**Figure 20.**
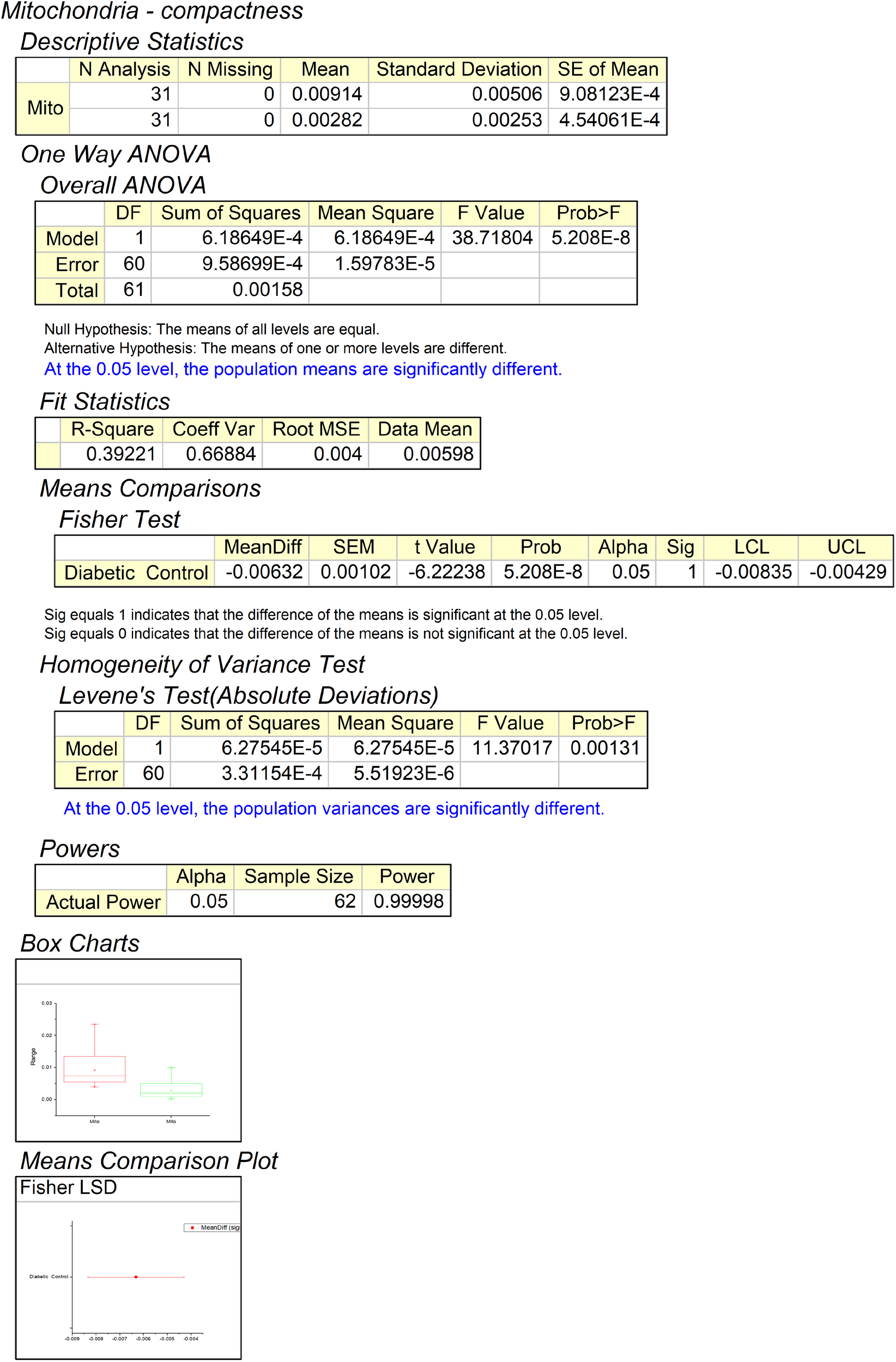
Hypothesis testing using one way ANOVA for mitochondria compactness significance between GAN outputs using control and diabetic dataset.

**Figure 21.**
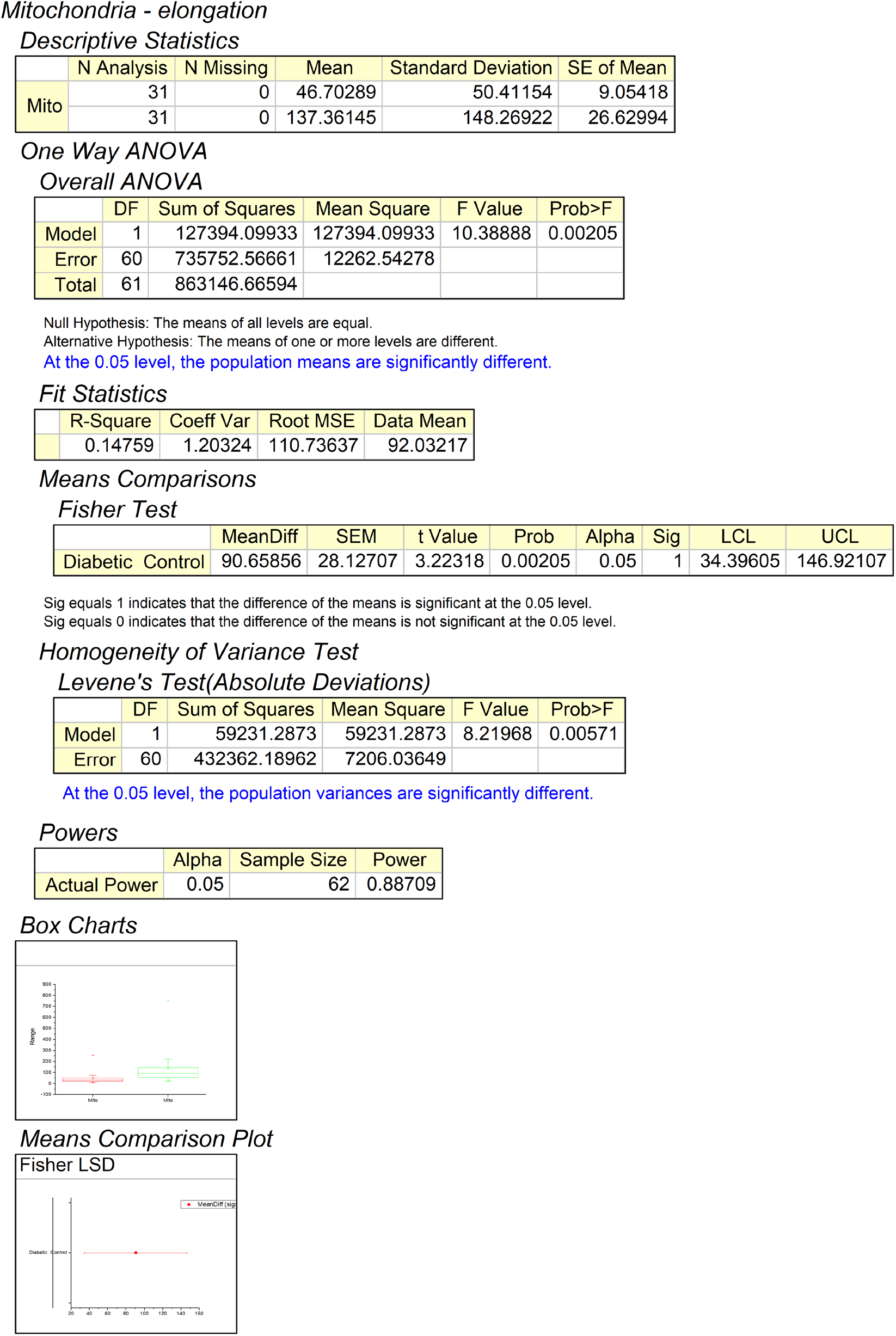
Hypothesis testing using one way ANOVA for mitochondria elongation significance between GAN outputs using control and diabetic dataset.

**Figure 22.**
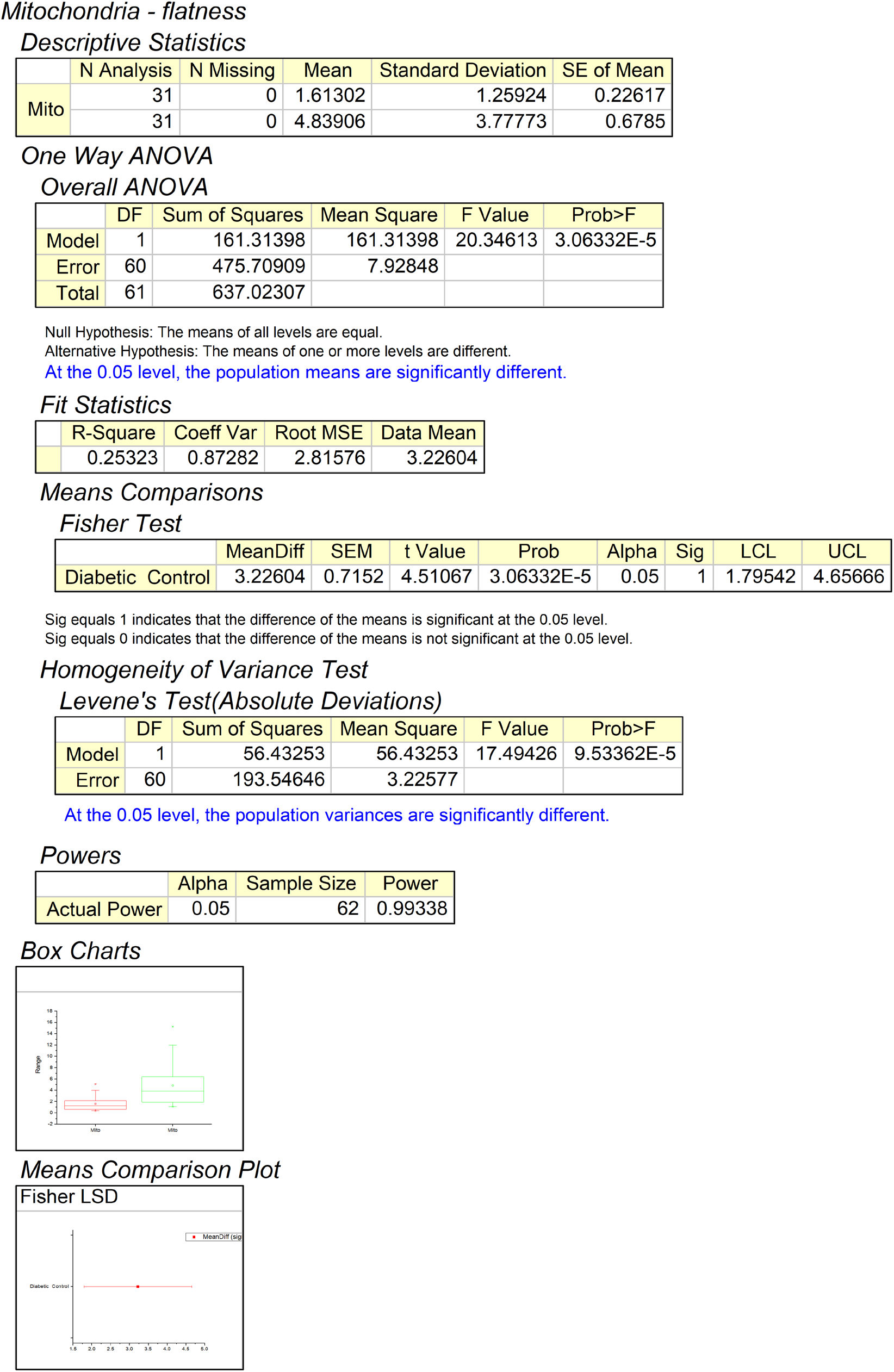
Hypothesis testing using one way ANOVA for mitochondria flatness significance between GAN outputs using control and diabetic dataset.

**Figure 23.**
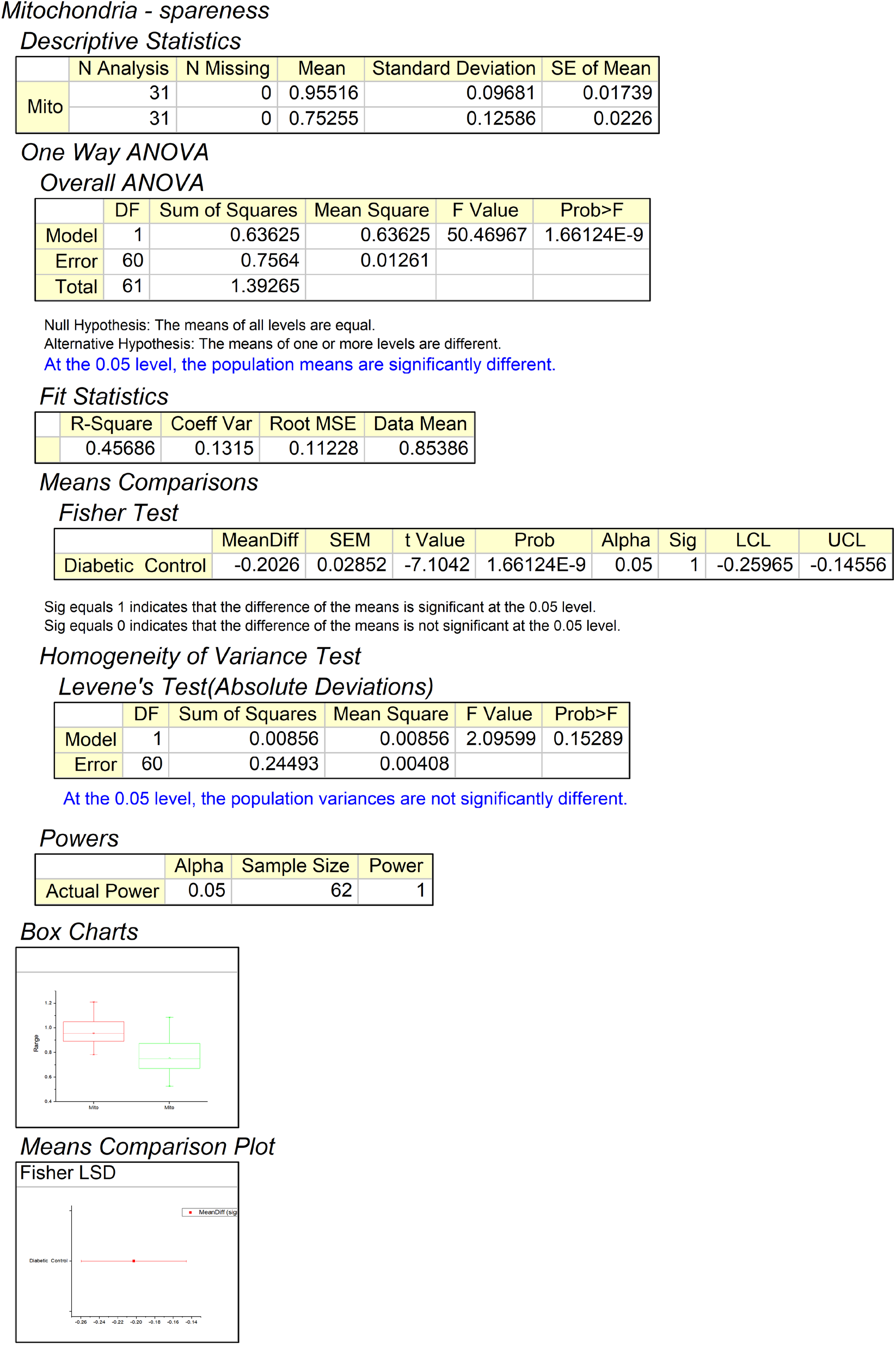
Hypothesis testing using one way ANOVA for mitochondria spareness significance between GAN outputs using control and diabetic dataset.

**Figure 24.**
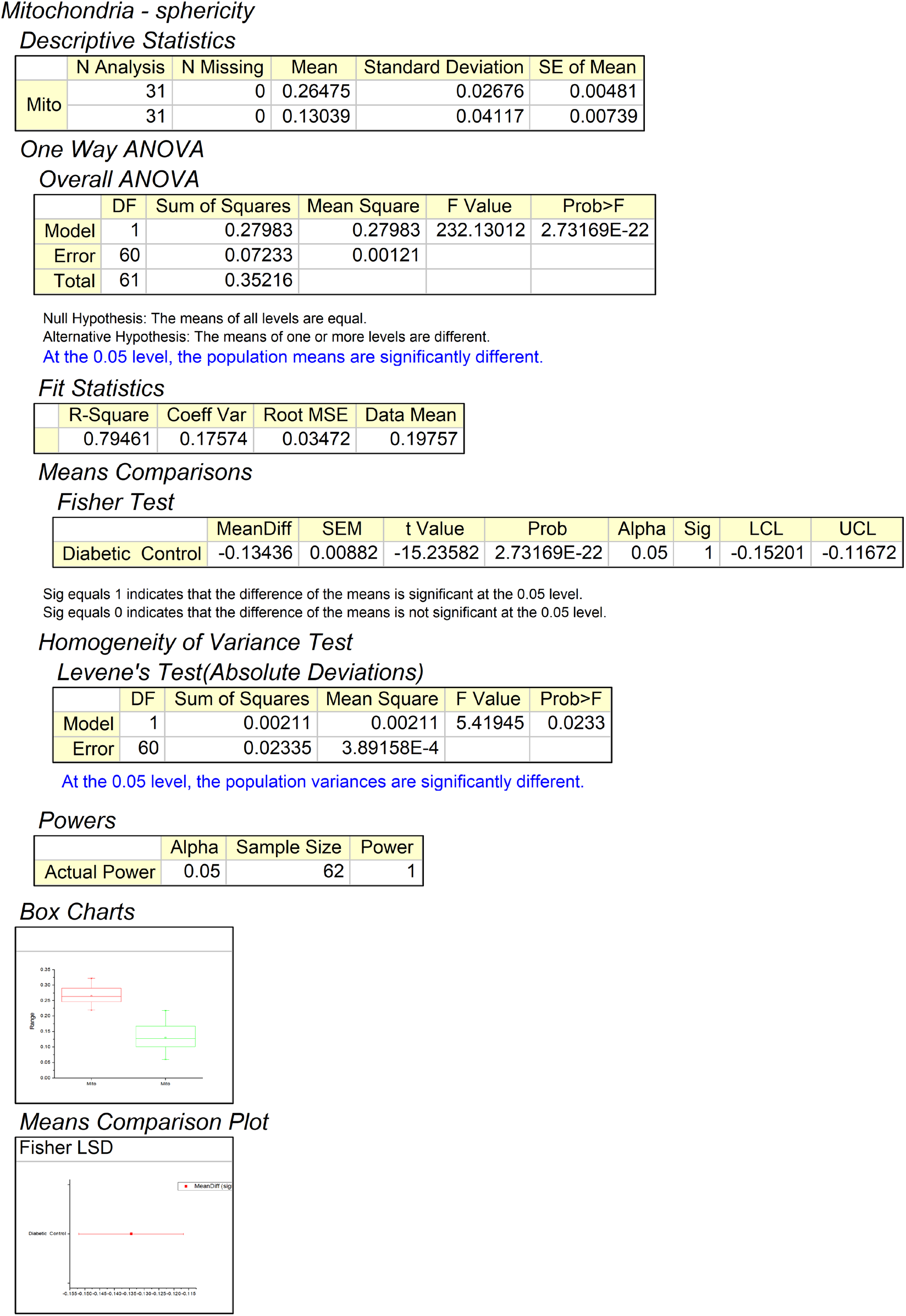
Hypothesis testing using one way ANOVA for mitochondria sphericity significance between GAN outputs using control and diabetic dataset.

**Figure 25.**
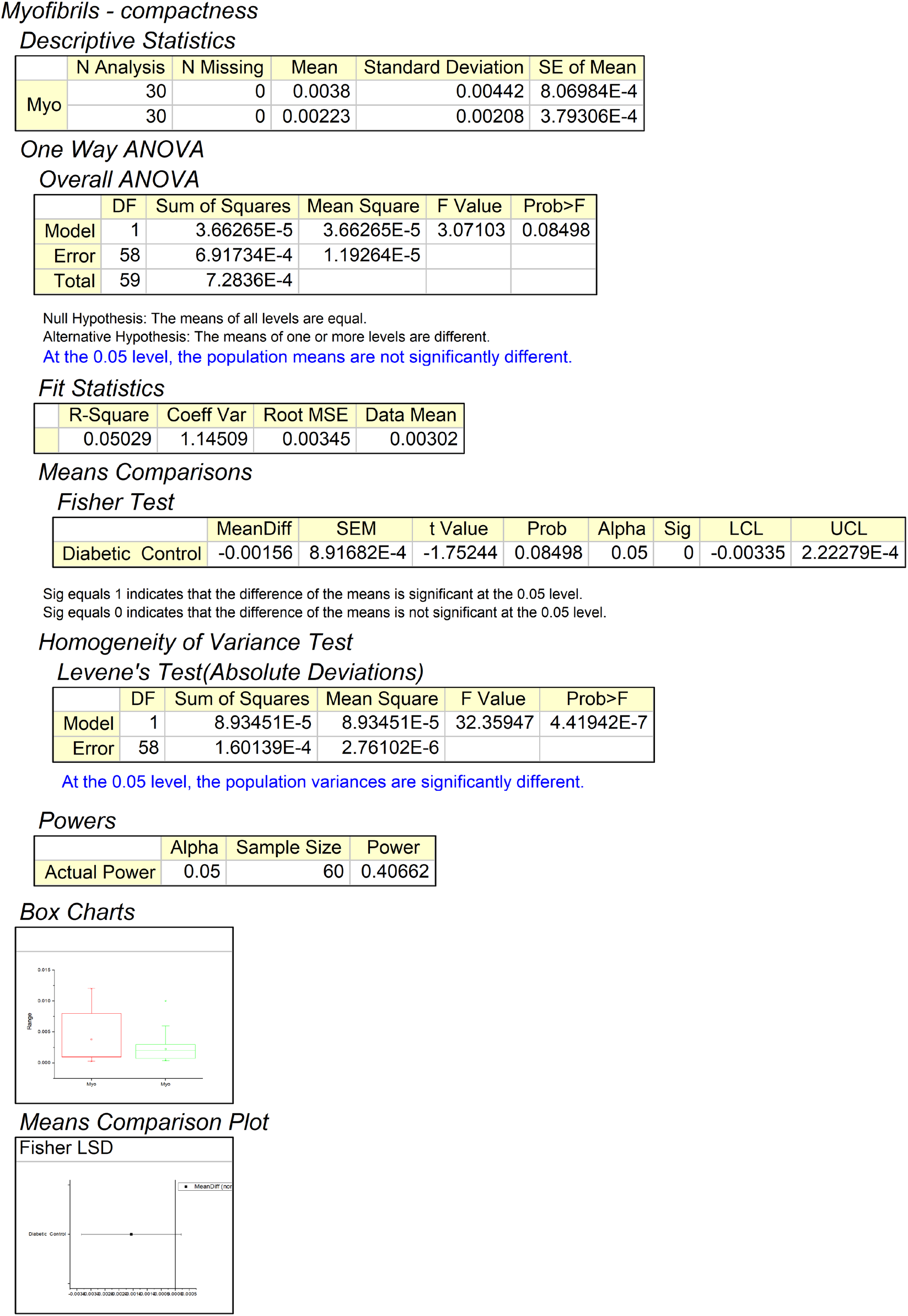
Hypothesis testing using one way ANOVA for myofibrils compactness significance between GAN outputs using control and diabetic dataset.

**Figure 26.**
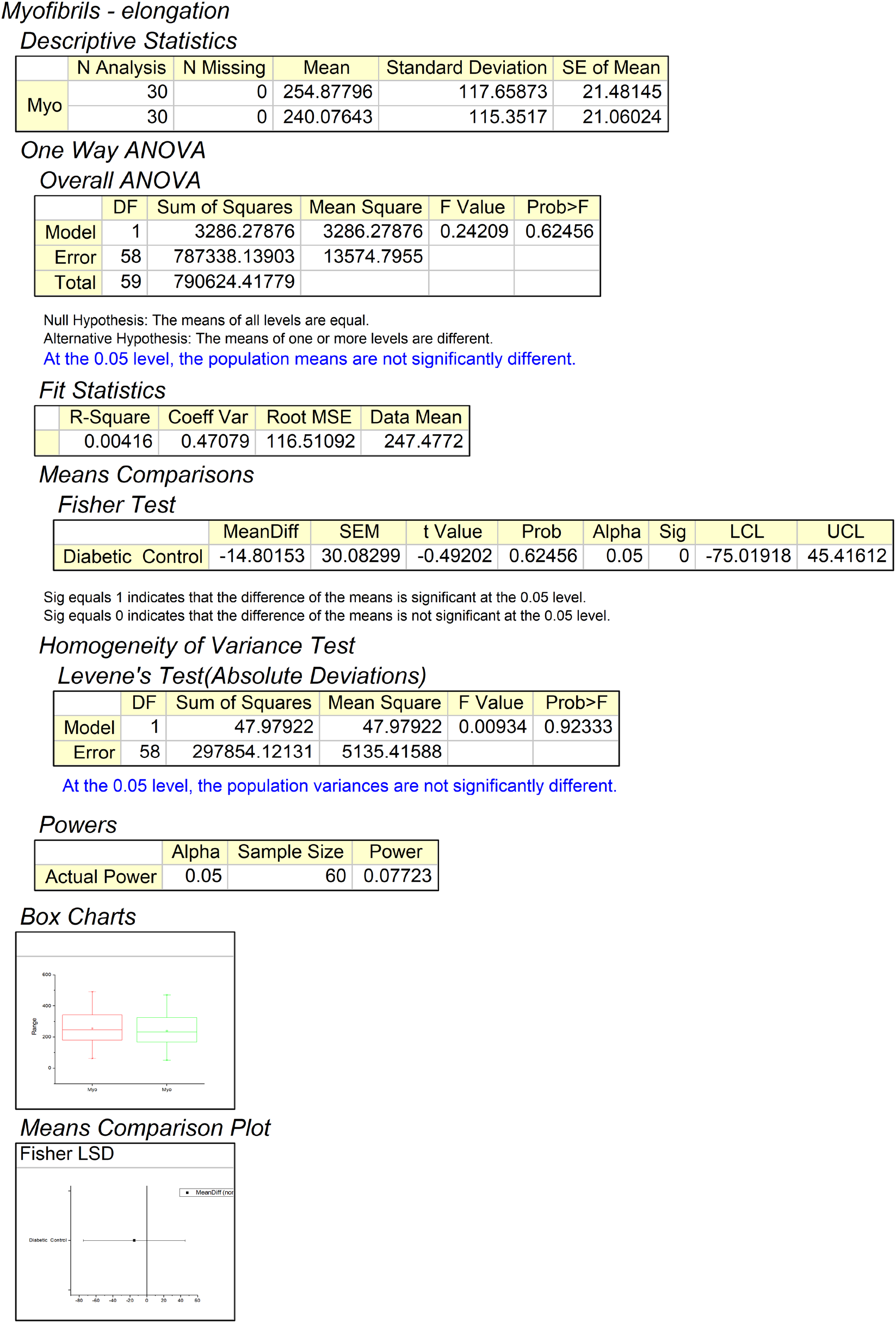
Hypothesis testing using one way ANOVA for myofibrils elongation significance between GAN outputs using control and diabetic dataset.

**Figure 27.**
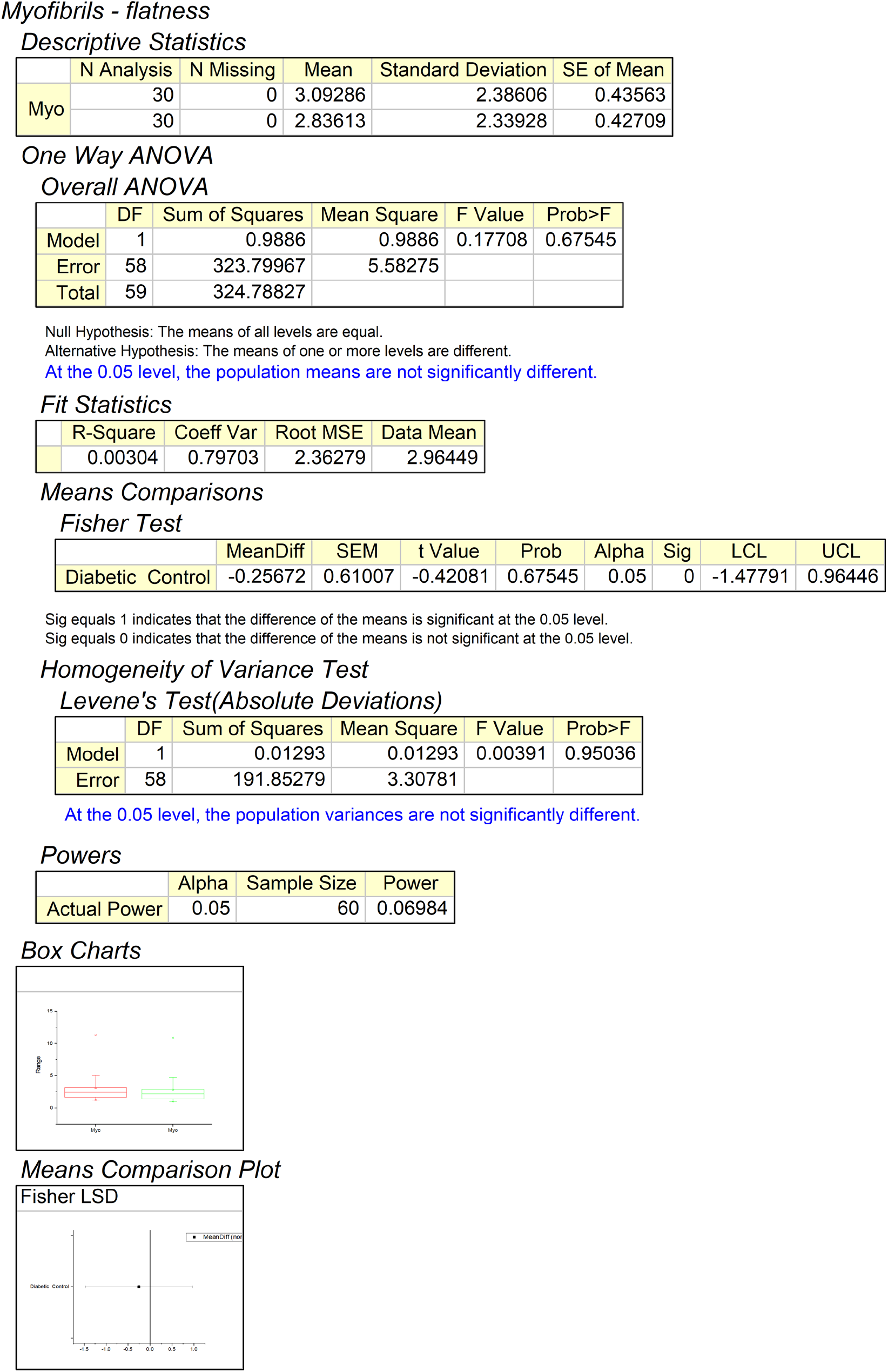
Hypothesis testing using one way ANOVA for myofibrils flatness significance between GAN outputs using control and diabetic dataset.

**Figure 28.**
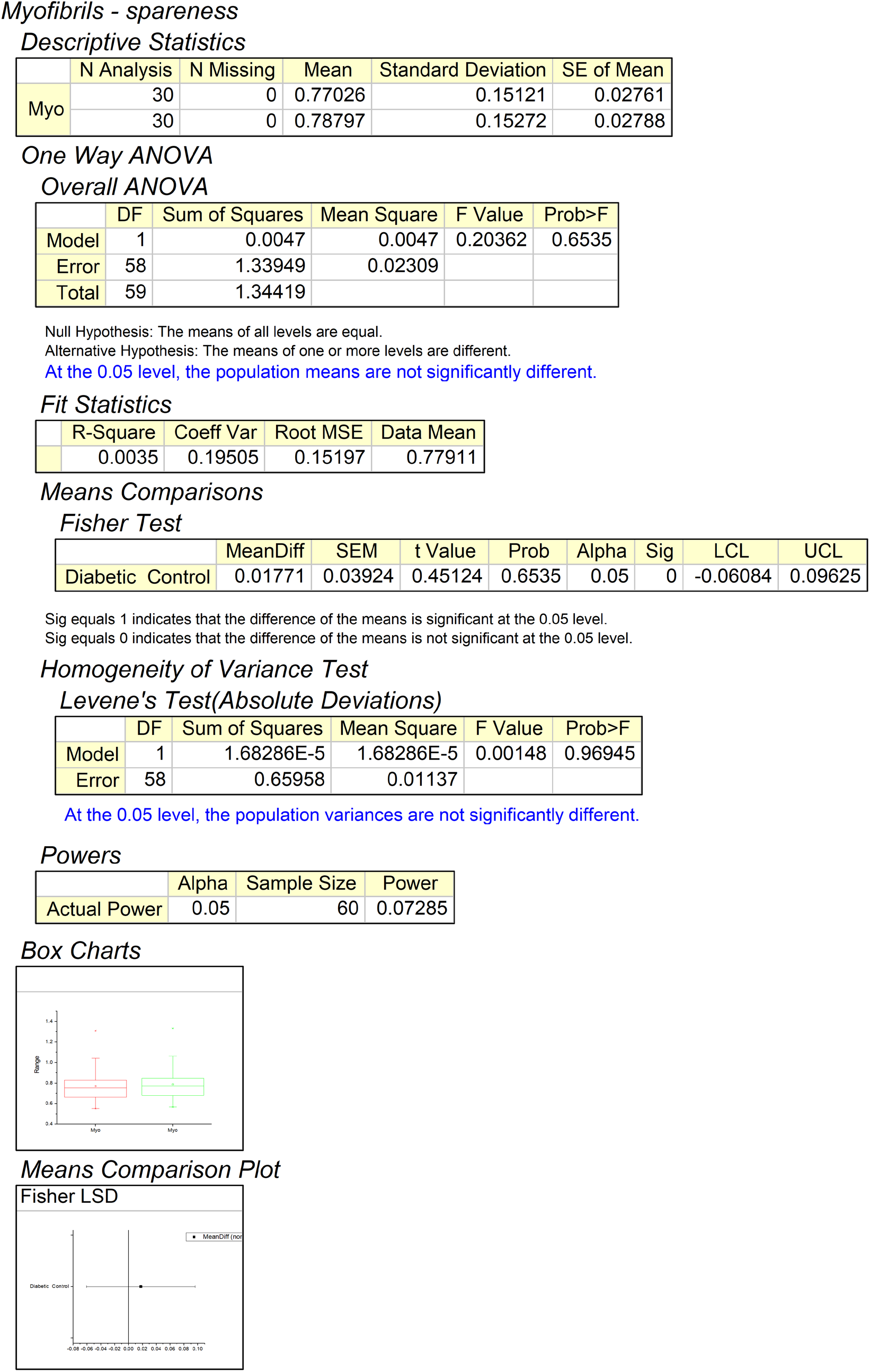
Hypothesis testing using one way ANOVA for myofibrils spareness significance between GAN outputs using control and diabetic dataset.

**Figure 29.**
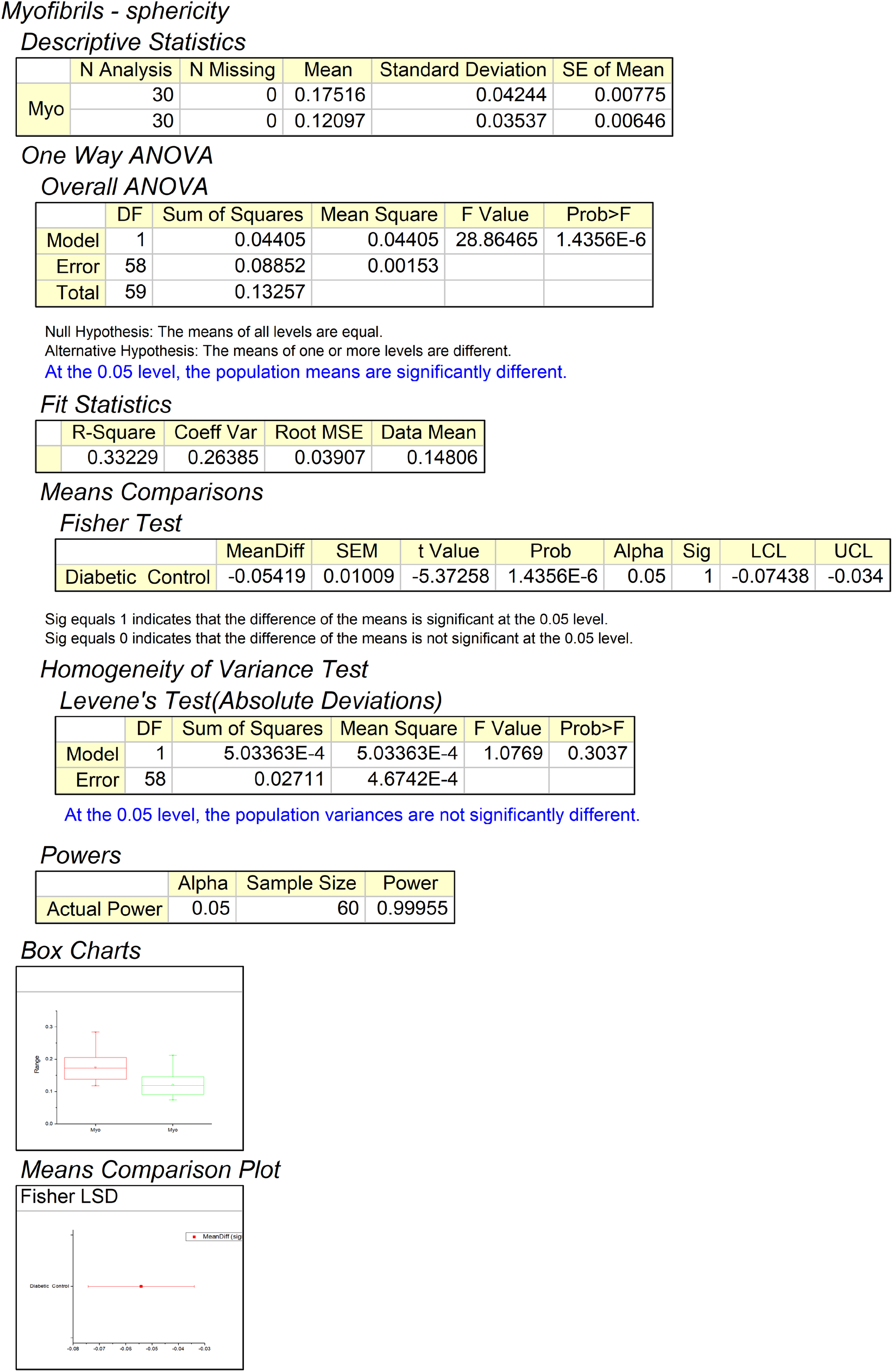
Hypothesis testing using one way ANOVA for myofibrils sphericity significance between GAN outputs using control and diabetic dataset.

## References

1. Khadangi, A., T. Boudier, and V. Rajagopal, EM-stellar: benchmarking deep learning for electron microscopy image segmentation. Bioinformatics, 2021. 37(1): p. 97–106.

2. Khadangi, A., T. Boudier, and V. Rajagopal. EM-net: Deep learning for electron microscopy image segmentation. in 2020 25th International Conference on Pattern Recognition (ICPR). 2021. IEEE.

3. Johnson, G.R., R.M. Donovan-Maiye, and M.M. Maleckar, Generative modeling with conditional autoencoders: Building an integrated cell. arXiv preprint arXiv:1705.00092, 2017.

4. Shaga Devan, K., et al., Improved automatic detection of herpesvirus secondary envelopment stages in electron microscopy by augmenting training data with synthetic labelled images generated by a generative adversarial network. Cellular Microbiology, 2021. 23(2): p. e13280.

5. Goldsborough, P., et al., CytoGAN: generative modeling of cell images. BioRxiv, 2017: p. 227645.

6. Lu, A.X., et al., Learning unsupervised feature representations for single cell microscopy images with paired cell inpainting. PLoS computational biology, 2019. 15(9): p. e1007348.

7. Yuan, H., et al., Computational modeling of cellular structures using conditional deep generative networks. Bioinformatics, 2019. 35(12): p. 2141–2149.

8. Phillip, J.M., et al., A robust unsupervised machine-learning method to quantify the morphological heterogeneity of cells and nuclei. Nature protocols, 2021. 16(2): p. 754–774.

9. Karras, T., S. Laine, and T. Aila. A style-based generator architecture for generative adversarial networks. in Proceedings of the IEEE/CVF Conference on Computer Vision and Pattern Recognition. 2019.

10. Majarian, T.D., et al., CellOrganizer: Learning and Using Cell Geometries for Spatial Cell Simulations. Methods in molecular biology (Clifton, NJ), 2019. 1945: p. 251.

11. Jarosz, J., et al., Changes in mitochondrial morphology and organization can enhance energy supply from mitochondrial oxidative phosphorylation in diabetic cardiomyopathy. American Journal of Physiology-Cell Physiology, 2017. 312(2): p. C190–C197.

12. Su, M., et al., Generative adversarial networks as a tool to recover structural information from cryo-electron microscopy data. BioRxiv, 2018: p. 256792.

13. Ghahramani, A., F.M. Watt, and N.M. Luscombe, Generative adversarial networks simulate gene expression and predict perturbations in single cells. BioRxiv, 2018: p. 262501.

14. Comes, M.C., et al., Multi-scale generative adversarial network for improved evaluation of cell–cell interactions observed in organ-on-chip experiments. Neural Computing and Applications, 2021. 33(8): p. 3671–3689.

15. Aida, S., et al., Deep Learning of Cancer Stem Cell Morphology Using Conditional Generative Adversarial Networks. Biomolecules, 2020. 10(6): p. 931.

16. Lopez, R., et al., Deep generative modeling for single-cell transcriptomics. Nature methods, 2018. 15(12): p. 1053–1058.

17. Khadangi, A., E. Hanssen, and V. Rajagopal. Automated framework to reconstruct 3D model of cardiac Z-disk: an image processing approach. in 2018 IEEE International Conference on Bioinformatics and Biomedicine (BIBM). 2018. IEEE.

18. Khadangi, A., E. Hanssen, and V. Rajagopal, Automated segmentation of cardiomyocyte Z-disks from high-throughput scanning electron microscopy data. BMC Medical Informatics and Decision Making, 2019. 19(6): p. 1–14.

19. Ronneberger, O., P. Fischer, and T. Brox. U-net: Convolutional networks for biomedical image segmentation. in International Conference on Medical image computing and computer-assisted intervention. 2015. Springer.

